# Structural basis of Rho-dependent transcription termination

**DOI:** 10.1101/2022.08.29.505764

**Authors:** Vadim Molodtsov, Chengyuan Wang, Emre Firlar, Jason T. Kaelber, Richard H. Ebright

## Abstract

Rho and NusG mediate factor-dependent transcription termination in *Escherichia coli*. Here, we report preparation of complexes functional in factor-dependent termination from RNA polymerase (RNAP), Rho, NusG, and synthetic nucleic-acid scaffolds, and we report cryo-EM structures of complexes. The structures show that functional factor-dependent pre-termination complexes contain a closed-ring Rho hexamer, have RNA threaded through the central channel of Rho, have 60 nt of RNA interacting sequence-specifically with the exterior of Rho and 6 nt of RNA interacting sequence-specifically with the central channel of Rho, have Rho oriented relative to RNAP such that ATP-hydrolysis-dependent translocation by Rho exerts mechanical force on RNAP, and have NusG bridging Rho and RNAP. The results explain five decades of research on Rho and provide a foundation for understanding Rho function.

**One sentence summary:** Cryo-EM reveals the structure of the functional Rho pre-termination complex

## Main Text

The ring-shaped hexameric ATP-dependent molecular motor Rho mediates factor-dependent transcription termination and transcription-translation-coupling quality control in *Escherichia coli* (reviewed in *1-4*). Rho is a close homolog of F_1_ of the F_1_F_0_ ATP synthase/ATPase, a ring-shaped hexameric molecular motor that employs a transmembrane proton gradient to drive synthesis of ATP, or, running in reverse, that employs ATP hydrolysis to drive generation of a transmembrane proton gradient (reviewed in *5*).

Five decades of genetic and biochemical experiments indicate that Rho-dependent transcription termination involves four steps (Fig. 1A; *1-4, 6-23*). In the first step, the Rho hexamer in an open-ring state binds to RNA emerging from a transcription elongation complex (TEC), recognizing a long C-rich RNA sequence (Rho utilization site, *rut* site) through a primary binding site (PBS) on the exterior of the Rho hexamer (Fig. 1A, first image). In the second step, the Rho hexamer in the open-ring state recognizes a short pyrimidine-rich RNA sequence through a secondary binding site (SBS) in the central channel of the Rho hexamer (Fig. 1A, second image). In the third step, the Rho hexamer transitions from the catalytically inactive, open-ring state to a catalytically competent, closed-ring state, with RNA threaded through the central channel of the Rho hexamer; this reaction is facilitated by interaction with transcription elongation factor NusG (Fig. 1A, third image). In the fourth step, the Rho hexamer in the catalytically competent, closed-ring state performs ATP-hydrolysis-dependent 5’→3’ translocation on RNA toward the TEC, exerting mechanical force on, and disrupting, the TEC (Fig. 1A, fourth image).

**Fig. 1.**
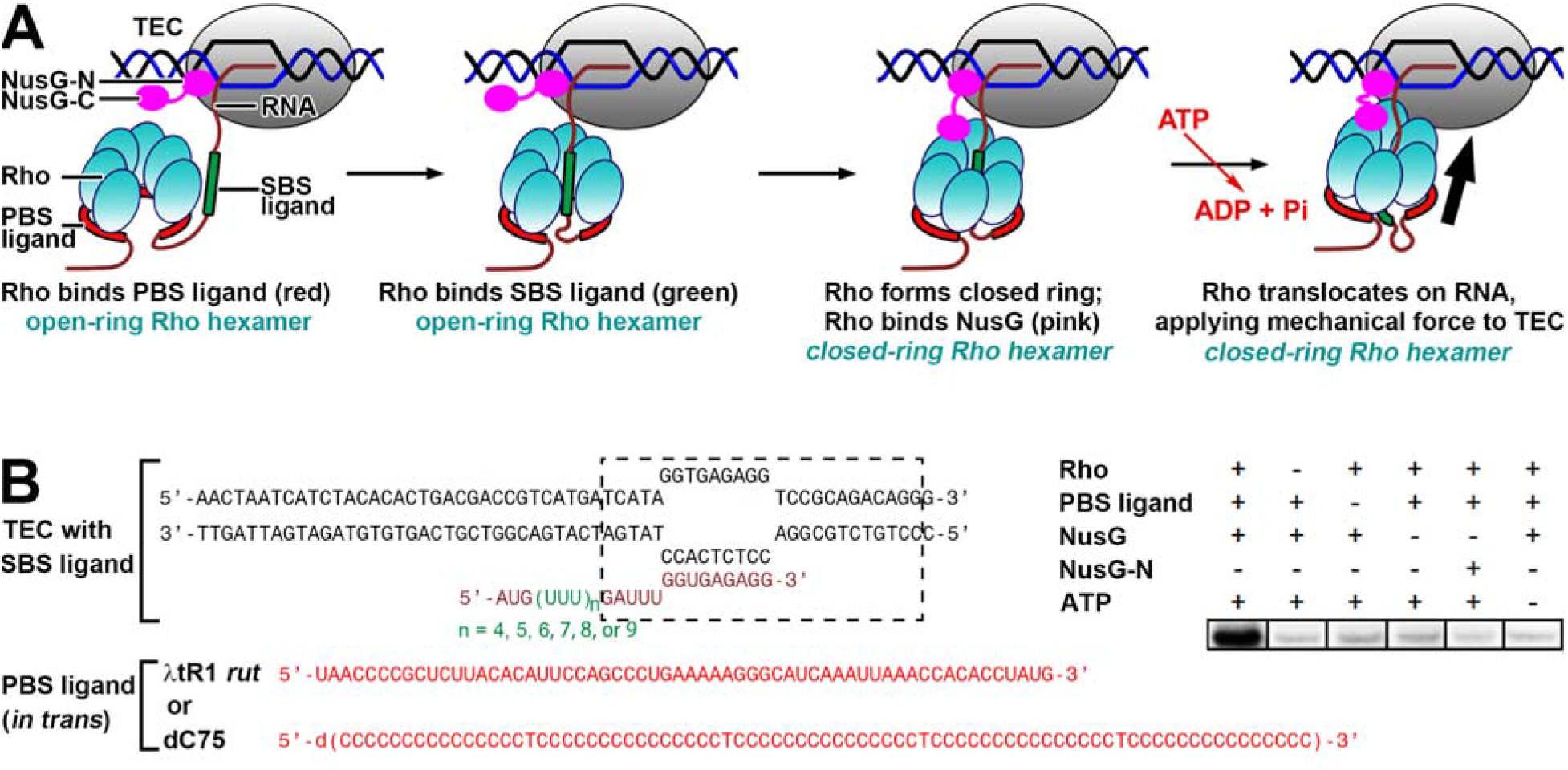
Rho-dependent termination. (**A)** Proposed mechanism of Rho-dependent termination (*1-4*). In the first step (image 1), the Rho hexamer in an open-ring state--either free or in complex with a TEC-- recognizes a long C-rich RNA sequence (PBS ligand; also known as *rut* site) through a primary binding site (PBS) on the exterior of the Rho hexamer (image 1). In the second step, the Rho hexamer in an open-ring state recognizes a short pyrimidine-rich RNA sequence (SBS ligand) through a secondary binding site (SBS) in the central channel of the Rho hexamer (image 2). In the third step, the Rho hexamer transitions from the catalytically inactive, open-ring state to the catalytically competent, closed-ring state; this reaction is facilitated by interactions with transcription elongation factor NusG (image 3). In the fourth step, the Rho hexamer in the catalytically active, closed-ring state performs ATP-hydrolysis-dependent 5’→3’ translocation on RNA toward the TEC, exerting mechanical force on, and disrupting, the TEC (image 4). Rho, cyan; RNAP in TEC, gray; nontemplate-strand DNA, template-strand DNA, and RNA, black, blue, and brick-red, respectively; PBS ligand, red; SBS ligand, green; RNA 5’ to PBS ligand, RNA between PBS ligand and SBS ligand, and spacer RNA between SBS ligand and TEC, brick-red; NusG, magenta. **(B)** Scaffold assay for Rho-dependent termination. Left, synthetic nucleic-acid scaffolds comprising: (i) determinants for formation of a TEC (nontemplate- and template-strand DNA oligomers in black, and RNA oligomer in brick-red and green; boundaries of TEC denoted by dashed rectangle); (ii) a 5’ extension of the RNA oligomer containing 4, 5, 6, 7, 8, or 9 codons of U-tract RNA as SBS ligand; and (iii) an RNA oligomer carrying the λtR1 *rut* site (red) or dC75 (*23*; red), provided *in trans*, as PBS ligand. Right, RNA-release data showing that scaffold with λtR1 *rut* site provided *in trans* is functional in Rho-dependent termination.

Four decades of genetic and biochemical research indicate that Rho mediates transcription-translation-coupling quality control by performing termination when, and only when, transcription-translation coupling is not occurring–for example, when the translation machinery stops synthesis but the TEC continues synthesis, translocating downstream and away from the translation machinery (*1-4, 25-30*).

Recently, two research teams reported cryo-EM structures of Rho-TEC complexes assembled on synthetic nucleic-acid scaffolds (*31-33*). However, the reported structures differed from expectation based on genetic and biochemical experiments in crucial respects (*4, 31-33*). The structures did not show the expected, catalytically competent, closed-ring state of Rho, did not show the expected protein-RNA interactions with the Rho PBS and Rho SBS, did not show the expected threading of RNA through Rho, did not show the expected orientation of Rho relative to the TEC, and did not show the expected interaction between Rho and NusG (*4, 31-33*). A particularly striking, functionally consequential, difference from expectation was the orientation of Rho relative to the TEC. In the structures, the open-ring Rho hexamer was oriented with its N-terminal PBS domains directed toward RNA polymerase (RNAP), and its C-terminal ATPase domains directed away from RNAP; this orientation is *opposite* to the expected orientation and is incompatible with ATP-hydrolysis-dependent 5’→3’ translocation on RNA toward the TEC.

The structures of *31* and *32* do not to correspond to any of the four functional states in the pathway of Rho-dependent termination expected based on the literature (Fig. 1A; *1-4, 31-33*). The authors argue that Rho-dependent termination proceeds through a mechanism that “dramatically deviates from the textbook paradigm” (*4, 31-33*): namely, a mechanism in which the ATP-hydrolysis-dependent molecular-motor activity of Rho is not essential (*4, 31-33*). However, the stark, essentially complete, discrepancy between their structures and the available genetic and biochemical results raises questions whether their structures represent *bona fide*, on-pathway, functionally relevant, states.

One specific basis for concern about the structures of *31* and *32* is that they were determined using complexes that had not been demonstrated to be functional in Rho-dependent termination. Rho-dependent \termination previously has been demonstrated for complexes obtained by use of *in vitro* or *in vivo* promoter-dependent transcription, but Rho-dependent termination has not previously been demonstrated for complexes, such as those of *31* and *32*, obtained by *in vitro* assembly on synthetic nucleic-acid scaffolds.

Here, we have developed nucleic-acid-scaffold assays for Rho-dependent transcription termination (Figs. 1B, S1-S2), we have used the assays to identify scaffold sequences and reaction conditions that enable assembly of Rho-dependent termination complexes functional in Rho-dependent termination (Figs. 1B, S1-S2), and, then, we employed those sequences and conditions to prepare complexes and to determine structures by use of cryo-EM (Figs. 2, S3-S5; Table S1). The structures obtained in this manner, in contrast to the structures of *31* and *32*, agree with--and account for, in detail--expectation from genetic and biochemical experiments (Figs. 2-4, S6-S12).

**Fig. 2.**
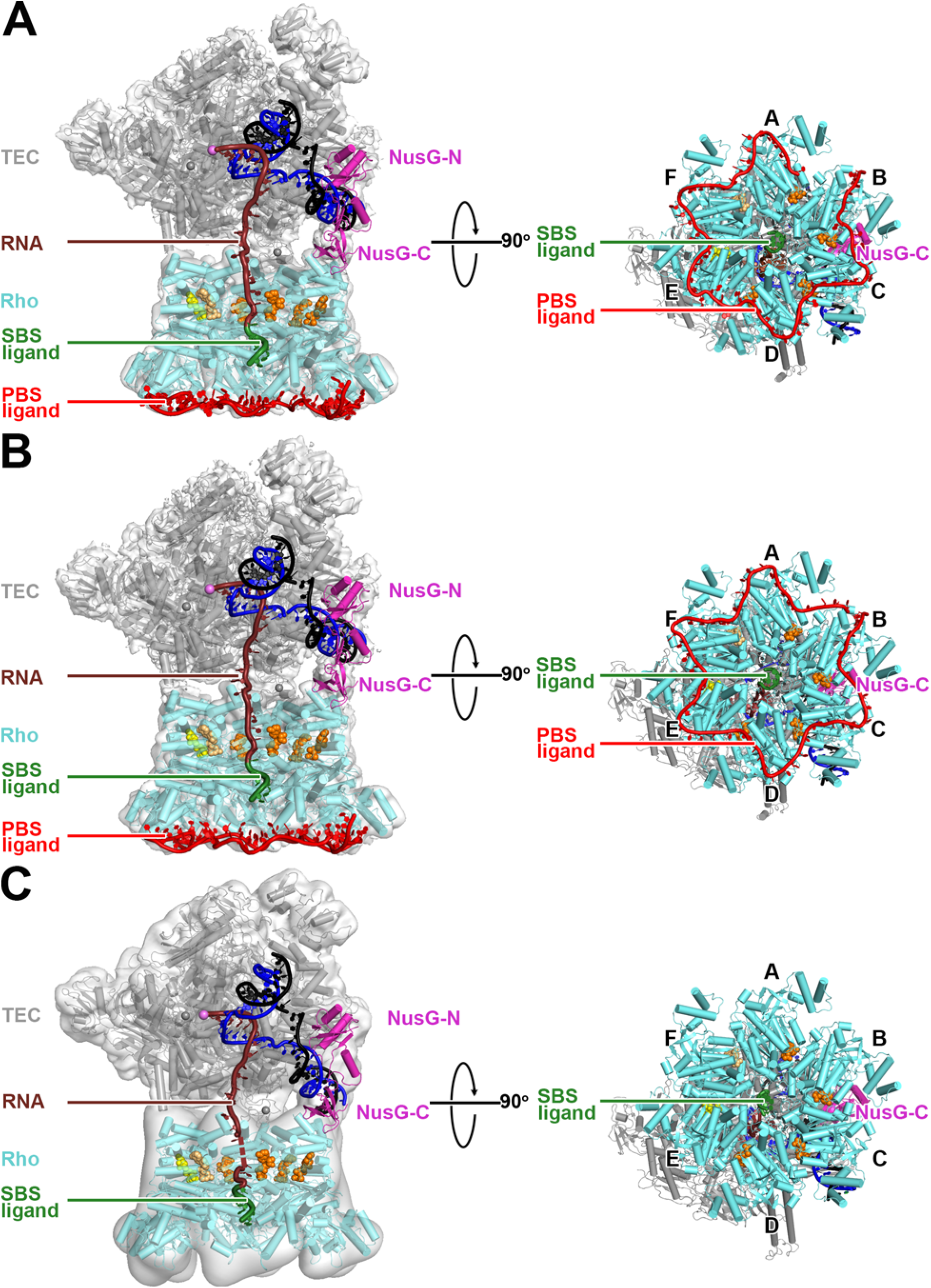
Structures of Rho pre-termination complexes. **(A)** Structure of complex with λtR1 *rut* RNA provided *in trans* as PBS ligand (λtR1-Rho-NusG-TEC). **(B)** Structure of complex with dC75 *rut* RNA provided *in trans* as PBS ligand (dC75-Rho-NusG-TEC). **(C)** Structure of complex with no PBS ligand (Rho-NusG-TEC). Left panels, view orientation having Rho-hexamer central channel aligned with y-axis, showing passage of RNA through TEC RNA-exit channel and through Rho-hexamer central channel. Right panels, orthogonal view orientation showing interaction of PBS ligand with Rho protomers A-F. Mg-ADP-BeF_3_ molecules, orange surfaces (high occupancy), light orange surfaces (low occupancy), and yellow surfaces (very low occupancy). Other colors as in Fig. 1A.

We prepared synthetic nucleic-acid scaffolds that comprised: (i) DNA and RNA determinants that direct formation of a TEC upon interaction with RNAP; (ii) a U-rich 5’ extension of the RNA, of length 4, 5, 6, 7, 8, or 9 codons, to serve as an SBS ligand and as a “spacer” between SBS ligand and TEC; and (iii) a second RNA oligomer, provided *in trans*, containing a PBS ligand, either the natural λtR1 *rut* site (*7*) or the synthetic mimic dC75 (*23*) (Fig. 1B, left). Using these scaffolds, we are able to demonstrate functional, efficient Rho-dependent transcription termination (Fig. 1B, right; Figs. S1-S2). The observed activity depends on Rho, depends on PBS ligand, depends on NusG--and cannot be substituted by NusG N-terminal domain (NusG-N), which interacts with RNAP but not with Rho (*1-4, 27-29*)--and depends on ATP (Fig. 1B, right). Both the λtR1 *rut* site provided *in trans* and dC75 provided *in trans* supported activity (Figs. 1, S1); shorter PBS ligands (dC5 and dC15) did not support efficient Rho-dependent termination, consistent with their previously demonstrated weaker interactions with Rho (Fig. S1; *22*). Equivalent results were obtained using 5’ U-rich extensions of the RNA of lengths 4, 5, 6, 7, 8, and 9 codons (Fig. S2A). Equivalent results were obtained using scaffolds having fully complementary DNA nontemplate and template strands (Fig. S1A) and nucleic-acid scaffolds having a partially noncomplementary sequence corresponding to the nontemplate strand of the transcription bubble (Fig. S1B). Equivalent results were obtained using RNA-release assays, in which Rho-dependent termination is assessed by detection of RNA released from immobilized TECs (Figs. 1, S1), and using RNA-extension assays, in which Rho-dependent termination is detected by cessation of RNA extension in the presence of all four NTPs (Fig. S2).

Having identified, in the nucleic-acid-scaffold assays, sequences and conditions that enable preparation of complexes proficient in Rho-dependent termination, we prepared complexes, we trapped complexes in a state poised to perform ATP-hydrolysis-dependent Rho translocation by replacing ATP by the ATP ground-state analog Mg-ADP-BeF_3_ (*23-24, 34*), and we determined structures of complexes by use of single-particle-reconstruction cryo-EM (Figs. 2, S2-S5). We determined three sets of structures: (i) a structure containing the λtR1 *rut* site (provided *in trans*) as PBS ligand, Rho, NusG, and TEC (λtR1-Rho-NusG-TEC; n = 6; 4.3 Å resolution), (ii) a structure containing dC75 (provided *in trans*) as PBS ligand, Rho, NusG, and TEC (dC75-Rho-NusG-TEC; 4.1 Å resolution), and (iii) structures containing only Rho, NusG, and TEC (Rho-NusG-TEC; 4.2-6.5 Å resolution) (Figs. 2, S3-S5).

The resulting structures differ radically from the structures of *31* and *32*. In λtR1-Rho-NusG-TEC and dC75-Rho-NusG-TEC, Rho adopts a catalytically competent, closed-ring state, superimposable on the catalytically competent, closed-ring state observed in crystal structures of Rho bound to Mg-ADP-BeF_3_ in the absence of a TEC (*23-24*; Fig. 2A-B), in contrast to the structures of *31* and *32*, in which Rho adopts a catalytically incompetent, open-ring conformation (Figs. 2 and S12). Comparison of λtR1-Rho-NusG-TEC and dC75-Rho-NusG-TEC structures to the structures of *30* and *31* indicates that the Rho hexamer is rotated by ∼150° about one axis, essentially flipping its orientation relative to the TEC, is rotated by ∼40° about a second axis, and is translated by ∼60-70 Å (Fig. S6). In λtR1-Rho-NusG-TEC and dC75-Rho-NusG-TEC, the central channel of the closed-ring Rho hexamer is aligned with the TEC RNA exit channel, and the closed-ring Rho hexamer is located immediately at the mouth of the TEC RNA exit channel, in a manner such that RNA proceeds directly from the TEC into, and through, the Rho hexamer (Fig. 2A-B), enabling interactions of the U-rich SBS ligand in the threaded RNA with the Rho SBS site (Fig. 2A-B), and interactions of the RNA oligomer, provided *in trans*, serving as PBS ligand, with the N-terminal domains of the six protomers of the Rho hexamer (Fig. 2A-B). In λtR1-Rho-NusG-TEC and dC75-Rho-NusG-TEC, Rho is oriented with its C-terminal ATPase domains toward the TEC and its N-terminal PBS domains away from the TEC, and thus, is in the orientation that enables ATP-hydrolysis-dependent Rho translocation toward the TEC (Fig. 2A-B). Crucially, in our structures, NusG bridges the TEC and Rho, with the NusG N-terminal domain (NusG-N) interacting with RNAP, and with the NusG C-terminal domain (NusG-C) interacting with the C-terminal domain of two protomers of Rho (Fig. 2A-B).

In the crystal structure of a catalytically competent, closed-ring state of Rho bound to an SBS ligand and Mg-ADP-BeF_3_ in the absence of NusG and TEC, the six Rho protomers exhibit small but clear differences in spacing, conformation, interactions, and occupancy levels of Mg-ADP-BeF_3_, corresponding to differences in motor state, and were assigned as protomers A-F (*4, 23*). In λtR1-Rho-NusG-TEC and dC75-Rho-NusG-TEC, the six Rho protomers exhibit the same small but clear differences in spacing, conformation, interactions, and occupancy levels of Mg-ADP-BeF_3_, and thus can be assigned in the same manner as protomers A-F (Fig. 2A-B, right). In λtR1-Rho-NusG-TEC and dC75-Rho-NusG-TEC, Rho is oriented relative to the TEC such that NusG-C interacts with the Rho protomers designated B and C (Fig. 2A-B, right).

The complex not having a PBS ligand, Rho-NusG-TEC, has the same overall structural organization as the complexes having a PBS ligand, λtR1-Rho-NusG-TEC and dC75-Rho-NusG-TEC (Fig. 2C vs. Fig. 2A-B; Fig. S5). This finding, together with our finding that the PBS ligand is required for Rho-dependent termination with our sequences and conditions (Figs. 1B, right, S1-S2), indicates that the PBS ligand is not essential for assembly of the Rho-NusG-TEC complex, but is essential for subsequent steps in Rho-dependent termination. We hypothesize that the PBS ligand facilitates subsequent steps in Rho-dependent termination allosterically, by inducing a conformational state or dynamic state of Rho competent for ATP-hydrolysis-dependent translocation. The lower map quality, especially for Rho, in our structures of complex not having a PBS ligand precludes defining details of the PBS-ligand-dependent change in conformation or dynamics, but is consistent with a PBS-ligand-dependent decrease in dynamics and increase in order of Rho.

In our structures of λtR1-Rho-NusG-TEC and dC75-Rho-NusG-TEC, we observe unambiguous, traceable density for 85-88 nt of RNA, comprising 14 nt interacting with the TEC, 7 nt in the spacer between the TEC and the Rho SBS, 6 nt of SBS-ligand RNA interacting with the Rho SBS, and 58-61 nt of PBS-ligand RNA interacting with the Rho PBS (58 nt in λtR1-Rho-NusG-TEC; 61 nt in dC75-Rho-NusG-TEC; Figs. 3, S3-S4). As such, the structures provide a *complete* description of protein-RNA interactions in the complexes.

**Fig. 3.**
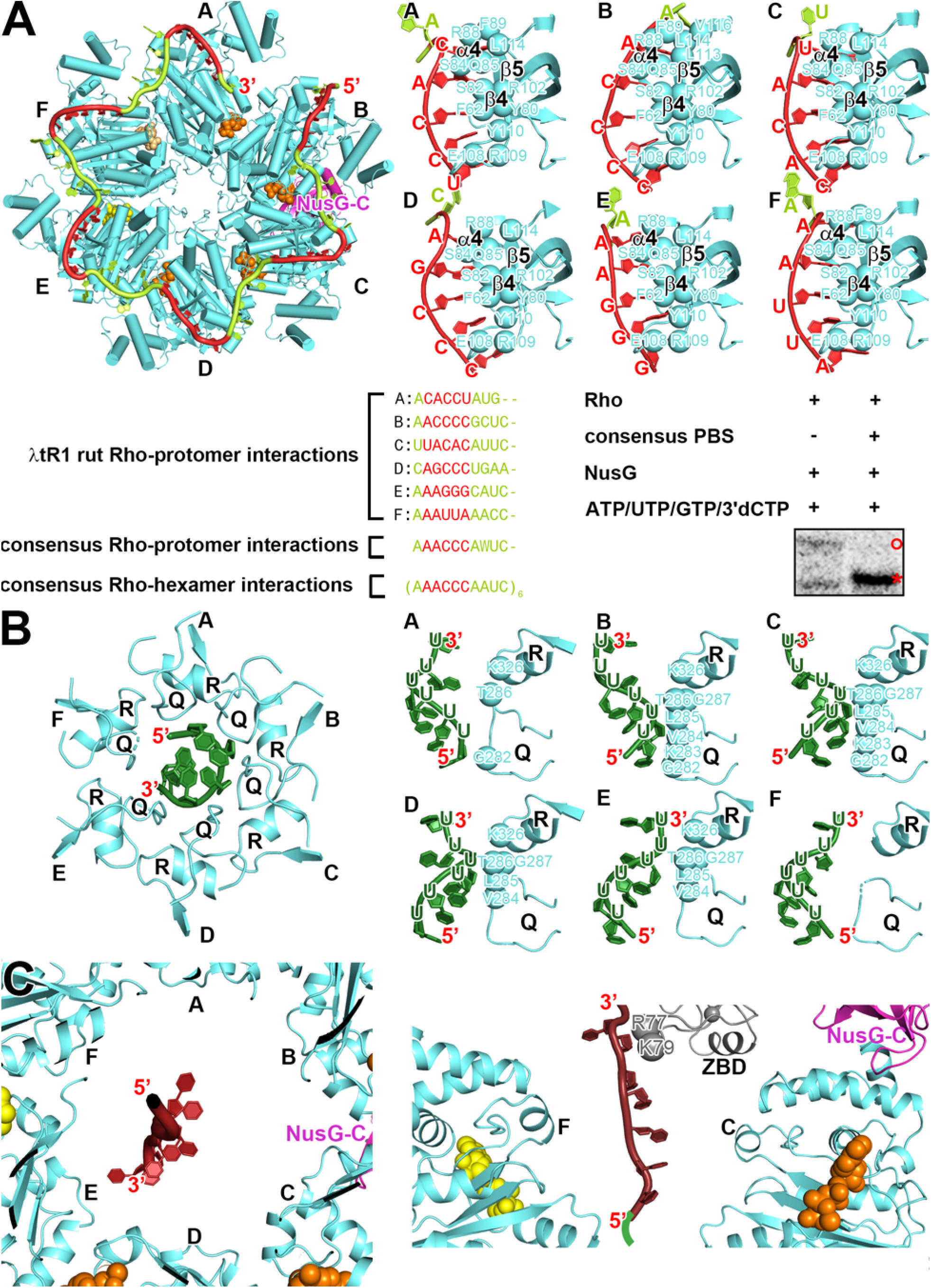
Protein-RNA interactions in Rho pre-termination complex. **(A)** Protein-RNA interactions by Rho PBS. Upper left, overall interaction (contacted nucleotides, red; non-contacted nucleotides, light green; view orientation and other colors as in Fig. 2, right). Upper right, details of interactions for Rho protomers A-F, Rho residues that interact with PBS ligand RNA are shown as cyan spheres. Lower left, 10 nt PBS-ligand sequences contacted by Rho protomers A, B, C, D, E, and F in λtR1-Rho-NusG-TEC, 10 nt inferred PBS-ligand consensus sequence for Rho protomer interaction, and 60 nt inferred PBS-ligand consensus sequence for Rho hexamer interaction, colors as in A. Lower right, RNA-extension data showing that an RNA oligomer carrying the inferred PBS-ligand consensus sequence is functional in Rho-dependent termination. **(B)** Protein-RNA interactions by Rho SBS. Left, overall interaction, focusing on 6 nt SBS ligand (green) and structural elements of Rho that interact with SBS ligand (Q-loop and R-loop) (view orientation and other colors as in Fig. 2, right). Right, details of interactions of Rho protomers A, B, C, D, E, and F. **(C)** Protein-RNA interactions with spacer RNA between Rho SBS and TEC. Left, overall interaction, showing large distances, and absence of contacts, between Rho and spacer RNA (view orientation and colors as in Fig. 2, right; SBS ligand and Rho structural elements that interact with SBS ligand omitted for clarity). Right, orthogonal view, showing large distances, and absence of contacts, between Rho (protomers A-B and D-E omitted for clarity) and RNA, and showing interactions of 2 nt of spacer RNA with RNAP *β*’-subunit ZBD.

The structures of λtR1-Rho-NusG-TEC and dC75-Rho-NusG-TEC reveal the full interactions of the PBS ligand with the Rho hexamer (Fig. 3A). All six protomers of the Rho hexamer interact with PBS-ligand RNA. Rho protomer B--one of the two protomers that interacts with NusG-C--interacts with the 5’ end of the PBS ligand, and Rho protomer A interacts with the 3’ end of the PBS ligand (Fig. 3A, top). Ten nucleotides of PBS-ligand RNA are associated with each Rho protomer; 5 nt of the PBS-ligand RNA make potentially sequence-specific interactions with α-helix 4 and β strands 4-5 of the Rho protomer (Fig 3A, top, red), and 5 nt of the PBS-ligand RNA serve to connect the sequence-specifically recognized RNA segment associated with one Rho protomer with the sequence-specifically recognized RNA segment of the Rho next protomer (Fig 3A, top, green). The observed interactions are essentially identical for λtR1-Rho-NusG-TEC and dC75-Rho-NusG-TEC (Figs. 3A, top and S7). The interactions by 2 nt of the 10 nt of PBS-ligand RNA associated with each Rho protomer are superimposable on the interactions observed in a crystal structure of having a Rho hexamer interacting with six copies of a 2 nt RNA oligomer in the absence of NusG and TEC (Fig. S8; *18*). The interactions of the other 8 nt of the 10 nt of PBS-ligand RNA associated with each Rho protomer were not previously known. In the structures of *31*, interactions between the PBS ligand and Rho were limited to just one of the six Rho protomers (Fig S9A), and in the structures of *32*, interactions between the PBS ligand and Rho were completely absent (Fig. S9B). Indeed, in the structures, of *31* and *32*, structural elements of RNAP and template-strand DNA make interactions with the Rho PBS of three to four Rho protomers, sterically precluding interactions of PBS-ligand RNA with those Rho protomers (Fig. S9).

Our finding that each Rho protomer interacts with 10 nt of the PBS-ligand RNA enables us to divide the PBS ligand into six 10 nt segments, to align the six 10 nt segments to derive a consensus PBS-ligand sequence for Rho protomer interaction (AAACCCAAUC; Fig. 3A, bottom left), and, by concatenating six copies of that sequence, to derive a consensus PBS-ligand sequence for Rho hexamer interaction ([AAACCCAAUC]_6_; Fig. 3A, bottom left). We have synthesized the consensus PBS ligand for Rho hexamer interaction, and we have shown that it supports efficient Rho-dependent transcription termination (Fig. 3A, bottom right), indicating that the consensus PBS ligand is a functional PBS ligand. Our consensus PBS-ligand sequence defines a structure-based functional length (60 nt) and a structure-based functional sequence ([AAACCCAAUC]_6_) for the PBS ligand, significantly advancing the understanding of the sequence requirements for a PBS ligand (previously essentially limited to the requirement for a ∼80 nt sequence rich in C and poor in G; *1-4, 10*).

Six nucleotides of SBS-ligand RNA make tight interactions with the central channel of the closed-ring Rho hexamer (Fig. 3B), in a manner superimposable on interactions observed in a crystal structure of a Rho hexamer interacting with SBS-ligand RNA in the absence of NusG and TEC (*23*). The 6 nt of SBS-ligand RNA form a right-handed helix (Fig. 3B). Interactions with this right-handed helix are made by the Q and R-loops of Rho protomers A-E, which protrude from the Rho C-terminal ATPase domain into the central cannel of the Rho hexamer, and which are arrayed in a right-handed spiral that tracks the right-handed helix formed by SBS-ligand RNA (Fig. 3B). Rho protomer A interacts with nucleotides at the 5’ end of the SBS ligand; Rho protomers B, C, D, and E interact with successive nucleotides of the SBS ligand ordered in the direction 5’→3’; and Rho protomer F makes no interaction with SDS ligand (Fig 3B). The two Rho protomers that interact with NusG-C, protomer B and protomer C, make the most extensive and most highly ordered interactions with SBS ligand, and the Rho protomer that makes the most extensive interaction with NusG-C, protomer C, interacts with the central two nucleotides of the 6 nt right-handed helix formed by the SBS ligand (Fig. 3B). The interactions between SBS-ligand RNA and the Rho SBS can accommodate pyrimidine nucleotides, but cannot accommodate purine nucleotides without steric clash, accounting for the known preference for pyrimidine-rich SBS ligands (*4, 23*).

RNA within the TEC RNA-exit channel connects to the SBS ligand through a 7 nt spacer RNA (Fig. 3C). The spacer RNA passes through a wide portion of the central channel of the Rho hexamer and makes no interactions with Rho (Fig. 3C). The 2 nt of the spacer RNA closest to the TEC RNA-exit channel interact with residues of the RNAP *β*’-subunit zinc-binding domain (ZBD; Fig. 3C), these interactions with the ZBD are distinctive and are not observed in structures of TECs without additional factors (*35*) or in structures of functional transcription-translation complexes (*36-37*). The absence of protein-RNA interactions with most nucleotides of the spacer RNA suggests there may be few, or no, constraints on sequence of the spacer RNA

In our work we used a discontinuous RNA, with the PBS ligand provided *in trans* and the remaining RNA provided *in cis*. Model building based on our structures indicates that the 3’ end of the PBS ligand could be connected to the 5’ end of the PBS ligand through an RNA connector having a minimum length of 10-11 nt. Model building further suggests that there is freedom for bulging, looping, and/or secondary-structure formation in the RNA connector between the 3’ end of the PBS ligand and the 5’ end of the SBS ligand, and thus suggests the connector also could accommodate longer lengths of RNA.

The interface between the NusG-containing TEC and Rho in λtR1-Rho-NusG-TEC and dC75-Rho-NusG-TEC involves three protein-protein interactions: a large protein-protein interaction by NusG-C (498 Å^2^ buried surface area), and smaller interactions between the RNAP *β*-subunit flap-tip helix (FTH; 245 Å^2^ buried surface area) and the RNAP *ω* subunit (158.Å^2^ buried surface area; Fig. 4A).

**Fig. 4.**
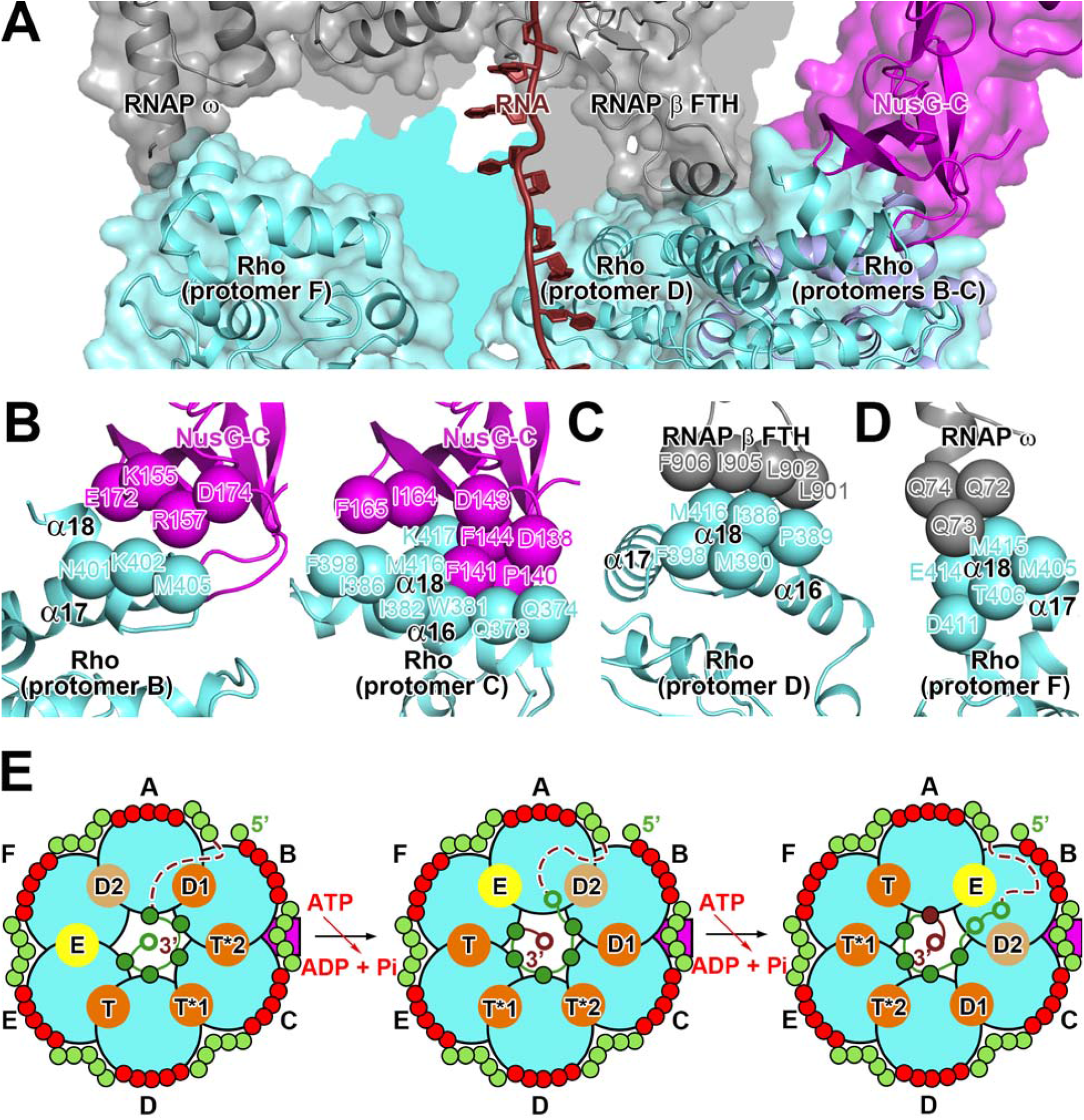
Rho-TEC interactions and ATP-binding-site motor states in Rho pre-termination complex. **(A)** Interface between Rho and NusG-containing TEC in λtR1-Rho-NusG-TEC (view orientation and colors as in Fig. 2, left). **(B)** Protein-protein interactions between Rho protomers B (left) and C (right) and NusG-C. **(C)** Protein-protein interactions between Rho protomer D and RNAP *β*-subunit FTH. **(D)** Protein-protein interactions between Rho protomer F and RNAP *ω* subunit. **(E)** Motor states of Rho ATP binding sites in Rho pre-termination complex (left), inferred motor states after first step of Rho-dependent ATP hydrolysis (center), and inferred motor states after second step of Rho-dependent ATP hydrolysis (right). Motor states are as follows: the ATP binding state (T), pre-catalysis ATP-bound state (T*1), catalytically competent ATP-bound state (T*2), ADP-bound state 1 (D1), ADP-bound state 2 (D2), and nucleotide exchange state (E) (*4, 23*). View orientation and colors as in Fig. 3A; PBS-ligand nucleotides, red circles for contact residues and light green circles for non-contact residues; SBS-ligand nucleotides, green circles (filled circles when interacting with SBS; open circles when not); spacer-RNA nucleotides, brick red circles (filled circles when interacting with SBS; open circles when not); orange, light orange, and yellow filled circles, Rho ATP binding sites (high-occupancy, low-occupancy, and very-low-occupancy ATP binding sites, respectively); magenta, NusG-C; curved black dashed lines, RNA connecting 3’ end of PBS ligand to 5’ end of SBS ligand.

In our structures, NusG bridges RNAP and Rho (Figs 2, 4A-B), NusG-N interacts with its previously defined binding site on RNAP (*38*), NusG-C interacts with Rho protomers B and C, and NusG-N and NusG-C are connected through a partly flexible linker (NusG linker; Figs. 2, 4A-B, and S10A). In our structures of λtR1-Rho-NusG-TEC and dC75-Rho-NusG-TEC there is unambiguous, traceable density for NusG-N, NusG-C, and all residues of the NusG linker (Figs. S3G, S4G). NusG-C wedges between the C-terminal ATPase domains of Rho protomer B and Rho protomer C (Figs 2, 4A-B, and S10A). NusG-C interacts extensively with the ATPase domain of Rho protomer C (388 Å^2^ buried surface area); the interaction involves the *β*7-*β*8 loop and *β*9-*β*10 loop of NusG and α-helices 16 and 18 of Rho protomer C (Figs. 4B, right, S10A). NusG makes a second, smaller interaction with the ATPase domain of Rho protomer B (110 Å^2^ buried surface area); this interaction involves the *β*8-*β*9 loop and *β*10-*β*11 loop of NusG-C and α-helix 17 of Rho protomer B (Figs. 4B, left, S10A). The interaction between NusG-C and Rho protomer C matches the interaction reported for a crystal structure of a Rho hexamer bound to six copies of a NusG-C protein fragment, with each of the six Rho protomers interacting with one copy of NusG-C protein fragment (Fig. S10B; *24*). The interaction between NusG-C and Rho protomer B was not previously described. It is not present in the crystal structure of a Rho hexamer bound to six copies of a NusG-C protein fragment (*24*), apparently because steric clash between adjacent NusG-C protein fragments in that structure resulted in disorder of the NusG *β*8-*β*9 loop and disruption of interactions made by the NusG *β*8-*β*9 and *β*10-*β*11 loops (Fig. S10B).

It previously has been shown that interaction of NusG-C with Rho facilitates formation of the catalytically competent, closed-ring state of the Rho hexamer (*1-4, 24*). The structures of λtR1-Rho-NusG-TEC and dC75-Rho-NusG-TEC reveal three additional functions of the interactions involving NusG. First, by bridging the TEC and Rho, NusG stabilizes the TEC-Rho interface (Figs. 2, 4A-B). In parallel structure-determination experiments performed in the presence and the absence of NusG, substantially lower numbers of particles of a size representing potential TEC-Rho complexes were observed in the absence of NusG than in its presence, providing direct support for the hypothesis that NusG stabilizes the TEC-Rho interface. Second, by interacting with Rho protomers B and C, NusG sets the rotational orientation of the Rho hexamer relative to the TEC (Figs. 2, 3, S10B), and thus sets the initial motor state of the Rho hexamer (Fig. 4E). One of the two Rho protomers that interacts with NusG, protomer B, serves as a binding site for the 5’ end of the PBS ligand and thus as the probable point of initial interaction between the PBS ligand and Rho (Fig. 3A); the two protomers of Rho that interact with NusG, protomers B and C, make the most highly ordered interactions with the SBS ligand (Fig. 3B, right); and the protomer of Rho that makes the most extensive interaction with NusG-C, protomer C, interacts with the crucial central position of the SBS ligand (Fig. 3B, right). The ATP binding site between the two protomers of Rho that interact with NusG, protomers B and C, is one of the two ATP binding sites exhibiting highest occupancy and highest order of the Mg-ADP-BeF_3_ ligand (Fig. S11) and exhibits binding-site occupancy, order, and geometry that indicate it is the ATP binding site at which ATP hydrolysis occurs first (Fig. 4E, see discussion below). Third, and likely most important, by wedging between Rho protomers B and C, NusG constrains the rotational orientation of the Rho hexamer relative to the TEC (Figs. 2, 3, S10B), preventing rotation of the Rho hexamer relative to the TEC during ATP-hydrolysis by Rho, thereby preventing futile cycles of ATP-hydrolysis-dependent rotation of Rho relative to the TEC and enforcing functional cycles of ATP-hydrolysis-dependent translocation of Rho relative to the TEC. NusG thus serves as a *stator* for the Rho-NusG-TEC motor complex, analogous, in all respects, to the stator formed by the *b* subunit of the F_1_F_0_ ATP synthase/ATPase motor complex (*5*).

The other protein-protein interactions at the Rho-TEC interface are: (i) an interaction between the RNAP *β*-subunit FTH and Rho protomer B, involving one face of the *β* FTH and α-helices 16 and 18 of Rho protomer B (Fig. 4C), and (ii) an interaction between the RNAP *ω* subunit and Rho protomer F, involving the C-terminal α-helix of *ω* and α-helices 17 and 18 of Rho protomer F (Fig. 4D). In the absence of NusG, or in the presence of a NusG derivative lacking NusG-C, Rho-dependent termination is dramatically reduced, but is not eliminated (Fig. S1; *1-4*). We infer that, in the absence of NusG, or in the presence of a NusG derivative lacking NusG-C, protein-protein interactions by the RNAP *β* FTH and RNAP *ω* partly, but not completely, constrain rotational orientation of Rho relative to the TEC during ATP-hydrolysis, and thus partly, but not completely, serve as the motor stator required for functional ATP-hydrolysis-dependent translocation of Rho relative to the TEC.

The ATP binding sites of Rho are located at the interfaces between successive pairs of Rho protomers (Figs. 2-4, S11). In a crystal structure of a catalytically competent, closed-ring Rho hexamer bound to Mg-ADP-BeF_3_ in the absence of NusG and the TEC, the six ATP binding sites are assigned--based on order, occupancy, and interactions made by the Mg-ADP-BeF_3_ ligand--as follows: a site for ATP loading (T), a site partly poised for ATP hydrolysis (T*1), a site fully poised for ATP hydrolysis (T*2), a first binding site for the ADP product of ATP hydrolysis (D1), a second binding site for the ADP product of ATP hydrolysis (D2), and a binding site for nucleotide release and exchange (E) (*4, 23*). In λtR1-Rho-NusG-TEC and dC75-Rho-NusG-TEC. the binding sites between Rho protomers D and E, C and D, B and C, A and B, A and F, and E and F correspond, respectively, to motor states T, T*1, T*2, D1, D2, and E (Figs. 4E, left, S11). The ATP binding site formed by the two Rho protomers that interact with NusG-C. protomers B and C, is the binding site in motor state T*2, and thus is the biding site at which ATP hydrolysis is predicted to occur first.

Our structures suggest the following, mechanism for ATP-hydrolysis-dependent translocation by Rho (Fig. 4E). In the first cycle of ATP-hydrolysis-dependent translocation of Rho, ATP hydrolysis occurs at the ATP binding site at the interface between two subunits contacted by NusG, protomers B and C (Fig 3E, left and center). This resets the motor state of the ATP binding site at the interface between protomers B and C from state T*2 to state D1, and increments the motor states of the other five ATP binding sites by one state (Fig. 4E, left vs. Fig. 4E, center). Concurrently, this results in relocation of the 5’ nucleotide of the SBS ligand out of the Rho SBS (Fig. 4E, left, green filled circle vs. Fig. 4E, center, green open circle), relocation of the 3’ nucleotide of the SBS ligand into the Rho SBS (Fig. 4E, left, green open circle vs. Fig. 4E, center, green filled circle), and relocation of the 5’ nucleotide of the RNA spacer into the position previously occupied by the 3’ nucleotide of the SBS ligand (Fig. 4E, center, brick-red open circle). The resulting translocation of Rho by 1 nt relative to the spacer RNA and the TEC applies mechanical force to the TEC, causing either forward translocation of the TEC by 1 nt without nucleotide addition (hypertranslocation model; *1, 3-4, 19*), extraction of RNA from the TEC by 1 nt (RNA-extraction model; *1, 3-4*), or reorganization of TEC structure (allosteric model; *1, 3-4, 20*), and thereby inhibits RNA extension by the TEC and predisposes the TEC to dissociation. In the second cycle of ATP hydrolysis, hydrolysis occurs at the ATP binding site at the interface between Rho protomers C and D. This converts the ATP binding site between Rho protomers C and D to state D1, increments the motor states of the other five ATP binding sites by one state (Fig. 4E, center vs. Fig. 4E, and right), and results in translocation of Rho by an additional 1 nt relative to the spacer RNA and the TEC, further applying mechanical force on the TEC, and thereby further inhibiting RNA extension by the TEC and further predisposing the TEC to dissociation (Fig. 4E, center and left). Subsequent cycles of ATP hydrolysis, carried out in the same manner, result in the further translocation by Rho relative to the spacer RNA and the TEC, further application of additional force to the TEC, and, ultimately, dissociation of the TEC.

Because the SBS ligand adopts the conformation of a right-handed helix (Fig. 3B), and because the Rho protomer Q and R-loops surround it in a right-handed spiral (Fig. 3B), translocation by Rho relative to RNA necessarily is accompanied by rotation of RNA within the central channel of the Rho hexamer. In principle, this rotation could apply a torque to the spacer RNA, the RNA in the TEC RNAP exit channel, and to the TEC RNA-DNA hybrid that could assist in unwinding of the upstream end of the RNA-DNA hybrid. However, in practice, our observation that a 5 nt segment of spacer RNA makes no protein-RNA interactions that could restrict rotation about RNA-backbone single bonds (Fig. 3C) suggests that torque may dissipate, in part or in whole, by rotation about RNA-backbone single bonds in spacer RNA, before reaching RNA in the TEC.

Our structures of λtR1-Rho-NusG-TEC and dC75-Rho-NusG-TEC agree, in all respects, with expectations from genetic and biochemical evidence results concerning Rho-dependent termination (*1-24*). The structures correspond to the third state (Fig. 1A, third subpanel) in what *32* and *33* refer to as the “textbook” model for Rho-dependent termination (Fig. 1A). The structures show the expected catalytically competent, closed-ring conformation of the Rho hexamer, show the expected threading of RNA through the central channel of the Rho hexamer, show the full PBS ligand interacting sequence-specifically with the full PBS of the Rho hexamer, show the full SBS ligand interacting sequence-specifically with the full SBS of the Rho hexamer, show Rho in the orientation relative to RNAP such that ATP-hydrolysis-dependent translocation can occur and can apply mechanical force to the TEC, and show NusG bridging the TEC and Rho (Figs. 2-4). We conclude that λtR1-Rho-NusG-TEC and dC75-Rho-NusG-TEC define the structure of the third state in the “textbook” model. the state in which the complex is fully assembled and fully poised to begin ATP-hydrolysis-dependent Rho translocation. We designate this state the “Rho pre-termination complex,” and we propose that it is the state functionally responsible for all or most Rho-dependent transcription termination in *E. coli*. As stated above, the structures of *31* and *32* lack these features and cannot be linked to any of the four states in the “textbook model” of Rho-dependent termination (Figs. 1A, S6). We suggest that the structures of *31* and *32* correspond to either: (i) on-pathway states that precede the four states in the “textbook” model in Fig. 1A (e.g., “pre-loading” states that precede full interaction of Rho with the PBS ligand), (ii) on-pathway states that have functional roles in Rho-dependent termination at certain sequences or under certain conditions, or (iii) off-pathway states that are non-functional and abiological.

Comparison of the structures of functional transcription-translation complexes (TTC-B; *30, 36-37*) and functional Rho pre-termination complexes (Figs. 2-4) establishes that the two sets of complexes are mutually exclusive (Fig. S12). The ribosome 30S subunit in TTC-B and Rho in the Rho pre-termination complex occupy the same face of the TEC; they cannot co-occupy without extensive steric overlap (Fig. S12). In addition, NusG bridges the ribosome 30S and the TEC in TTC-B, but bridges Rho and the TEC in the Rho pre-termination complex (Figs. S12). Comparison of the structures supports the following simple mechanism for Rho-dependent transcription-translation-coupling quality control (see *1-4, 27-30*): When transcription-translation coupling is occurring, Rho is unable to form a Rho pre-termination complex and unable terminate transcription; in contrast, when transcription-translation coupling is disrupted--for example, when the ribosome stalls but the TEC continues translocating downstream--Rho is able to form a Rho pre-termination complex and is able to terminate transcription. In this mechanism, both the TEC and NusG switch partners in the transition from transcription-translation-coupling to Rho-dependent termination. In transcription-translation coupling, the TEC makes protein-protein interactions with the ribosome, and NusG makes extensive interactions, mediated by NusG-C, with the ribosome (*1-4, 27-30, 36-37*). In contrast, in the Rho pre-termination complex, as discussed above, the TEC makes protein-protein interactions with Rho, and NusG makes extensive protein-protein interaction, mediated by NusG-C, with Rho (Fig. 4A-D).

The structures presented in this work restore the “textbook” model for Rho-dependent termination, necessitate reassessment of the functional roles, if any, of the structures of Rho-TEC complexes in *31* and *32*, and reveal, comprehensively, the protein-RNA interactions, protein-protein interactions, and protein-motor properties responsible of functional Rho pre-termination complexes. Key priorities for future work include determination of a structure of a Rho pre-termination complex containing a continuous RNA extending from the PBS ligand to the RNA 3’ end (as opposed to the current complexes, in which the PBS ligand is provided *in trans*; Figs. 1B-3), determination of a structures of a complex having the consensus PBS-ligand sequence defined in this work (Fig. 3A), determination of a structure of a complex that includes the transcription elongation factor NusA, which can be present in both TECs and TTCs (*1-4, 30, 37, 39*), and determination of structures of trapped intermediates after one, two, or more cycles of ATP-hydrolysis-dependent Rho translocation (Fig. 4E). The structures of this work thus provide an essential foundation for understanding Rho-dependent transcription termination and a platform for further structure determination illuminating the final, ATP-hydrolysis-dependent steps in Rho-dependent transcription termination.

## Acknowledgments

We thank Drs. James Berger and Irina Artsimovitch for plasmids and advice. We also thank the Rutgers CryoEM and Nanoimaging Facility, the National Center for CryoEM Access and Training (supported by NIH grant GM129539, Simons Foundation grant SF349247, and New York state grants), and the Stanford-SLAC Cryo-EM Center (supported by NIH grant GM129541) for microscope access.

## Author Contributions

V.M. and R.H.E. designed experiments. V.M., C.W., E.F., and J.T.K. collected data. V.M., C.W., and R.H.E. analyzed data. C.W. and R.H.E. prepared figures. R.H.E. wrote the manuscript.

## Data Availability Statement

Cryo-EM maps have been deposited in the Electron Microscopy Database (EMDB accession codes EMD-27928, EMD-27929, EMD-27930, EMD-27931, EMD-27932, EMD-27933, EMD-27864, EMD-27865, EMD-27897, EMD-27913, EMD-27914, EMD-27915, EMD-27916, EMD-27917, and EMD-27918), and atomic coordinates have been deposited in the Protein Database (PDB accession codes 8E3F, 8E3H, 8E5K, 8E5L, 8E5O, 8E5P, 8E6W, 8E6X, 8E6Z, and 8E70). Unique biological materials will be made available to qualified investigators on request.

## Supplementary Materials

## Materials and Methods

### *E. coli* RNAP core enzymes

*E. coli* RNAP core enzyme was prepared from *E. coli* strain BL21 Star (DE3) (ThermoFisher) transformed with plasmid pVS10 (*1*; encodes *E. coli* RNAP β’ with C-terminal hexahistidine tag, β, α, and ω subunits), as described (*2*?). The product (purity >99%) was stored in aliquots in RNAP storage buffer (10 mM Tris-HCl, pH 7.6, 100 mM NaCl, 0.1 mM EDTA, and 5 mM dithiothreitol) at -80°C.

### *E. coli* NusG

*E. coli* NusG was prepared from *E. coli* strain BL21 Star (DE3) (ThermoFisher) transformed with plasmid pIA247 (*3*; encodes *E. coli* NusG with C-terminal hexahistidine tag), as described (*4*). The product (purity >95%) was stored in aliquots in RNAP storage buffer at -80°C.

*E. coli* NusG-N (NusG^1-123^) was prepared from *E. coli* strain BL21 Star (DE3) (ThermoFisher) transformed with plasmid pRM431-NusG-N [constructed by site-directed mutagenesis of plasmid pRM431 (*5*; encodes *E. coli* NusG with N-terminal hexahistidine tag), using Vent DNA polymerase (New England Biolabs) and primers 5’-CAGCAGGTTGGTGATAAGCCGCGTTAGAAAACGCTGTTTGAACC GGGTGAAATGG-3’ and 5’-CCATTTCACCCGGTTCAAACAGCGTTTTCTAACGCGGCTTATCACC AACCTGCTG-3’] as described for NusG. The product (purity >95%) was stored in aliquots in RNAP storage buffer at -80°C.

### *E. coli* Rho

*E. coli* Rho was prepared from *E. coli* strain BL21 Star (DE3) (ThermoFisher) transformed with plasmid pET24b-Rho (encodes *E. coli* Rho; *6*). Cultures of transformants in 3 L LB broth (*7*) containing 30 *μ*g/ml kanamycin were shaken at 37°C until OD_600_ = 0.7, were induced by addition of IPTG and additional shaking for 3 h to at 37°C 1 mM. Cells were harvested by centrifugation 10 min at 5,000xg at 4°C, re-suspended in lysis buffer [20 mM Tris-HCl, pH 7.6, 50 mM NaCl, 1 mM EDTA, 1 mM β-mercaptoethanol, and EDTA-free protease inhibitor cocktail (Sigma-Aldrich), and lysed by sonication [S3000 sonicator (Misonix); 5 min; 5 s pulses at 50% maximum output; 2 s pauses between pulses] on ice. The cell lysate was cleared by centrifugation 30 min at 20,000xg at 4°C, the cleared lysate was applied to a 5 ml HiTrap Q HP column (GE Healthcare) equilibrated with TGE (*7*) containing 50 mM NaCl and 1 mM β-mercaptoethanol, and the column was eluted with a gradient of 50-500 mM NaCl in 30 column volumes of the same buffer. Pooled fractions containing Rho (∼300-400 mM NaCl) were concentrated to 2 ml by centrifugation 30 min at 4,000xg at 4°C through a 20 ml VivaSpin 20 30 kDa MWCO concentrator (Sartorius, and were diluted with TGE containing 1 mM β-mercaptoethanol to adjust the NaCl concentration to 50 mM. The resulting sample was applied to a 5 ml HiTrap HP heparin column (GE Healthcare) equilibrated with TGE containing 50 mM NaCl and 1 mM β-mercaptoethanol, and the column was eluted with a gradient of 50-700 mM NaCl in 40 column volumes of the same buffer. Pooled fractions containing Rho (∼300-400 mM NaCl) were concentrated to 2 ml by centrifugation 30 min at 4,000xg at 4°C through a VivaSpin 20 30 kDa MWCO concentrator. The resulting sample was applied to a 120 ml HiLoad 16/60 Superdex 200 size-exclusion column (GE Healthcare) equilibrated with 10 mM Tris-HCl, pH 7.6, 100 mM NaCl, and 1 mM β-mercaptoethanol, and the column was eluted with one column volume of the same buffer. Fractions containing pure Rho were pooled and concentrated to 100 *μ*M using a VivaSpin20 30 kDa MWCO concentrator. The product (purity >98%) was stored in aliquots at -80°C.

### Nucleic acids

Oligodeoxyribonucleotides and oligoribonucleotides (sequences in Figs. 1B, S1-S2 were purchased (Integrated DNA Technologies; PAGE-purified; nontemplate-strand oligodeoxyribonucleotides purchased in both non-biotinylated and 3’-biotinylated form), dissolved to 1 mM in 5 mM Tris-HCl, pH 7.5, and stored in aliquots at -80°C. Nucleic-acid scaffolds comprising nontemplate-strand DNA, template-strand DNA, and RNA product (sequences in Figs. 1B, S1-S2) were prepared by mixing 60 μM nontemplate-strand oligodeoxyribonucleotide, 60 μM template-strand oligodeoxyribonucleotide, and 60 μM RNA-product oligoribonucleotide or 5’-^32^P-labelled oligoribonucleotide [0.6 Bq/fmol; prepared by labelling of oligodeoxyribonucleotide using T4 polynucleotide kinase (New England BioLabs) and ATP or [*γ*^32^P]-ATP (Perkin-Elmer) in 100 µl annealing buffer (5 mM Tris-HCl, pH 7.5), heating 10 min at 95°C, and cooling over 3 h to 22°C; the resulting nucleic-acid scaffolds were stored in aliquots at -80°C. Partial nucleic-acid scaffolds comprising template-strand DNA and RNA product were prepared in the same manner, but omitting nontemplate-strand oligodeoxyribonucleotide.

### ADP-BeF_3_

ADP-BeF_3_ was prepared by reaction of 70 mM BeSO_4_ (Sigma-Aldrich) and 350 mM NaF (Sigma-Aldrich) in 50 µl 2 mM Tris-HCl, pH 7.0 for 1 min at 22°C, followed by addition of 15 *μ*l 100 mM ADP (Sigma-Aldrich). ADP-BeF_3_ was prepared freshly for each sample and was kept on ice until used.

### Scaffold assay for Rho-dependent transcription termination: RNA release

Reaction mixtures containing 0.5 µM RNAP core enzyme and 0.5 *μ*M partial nucleic-acid scaffold comprising template-strand oligodeoxyribonucleotide and 5’-^32^P-labelled RNA-product oligoribonucleotide (0.6 Bq/fmol) in 10 *μ*l transcription buffer (20 mM Tris-HCl, pH 7.9, 10 mM MgCl_2_, 20 mM KCl, and 1 mM DTT) were incubated 10 min at 22°C, were supplemented with 1 *μ*l 6 µM 3’-biotinylated nontemplate-strand oligodeoxyribonucleotide and incubated 5 min at 22°C, were supplemented with 1 *μ*l 0 or 30 µM NusG derivative (NusG or NusG-N) and incubated 5 min at 22°C, were supplemented with PBS-ligand oligoribonucleotide (1 *μ*l 0 or 50 *μ*M λtR1 *rut*, 1 *μ*l 50 *μ*M dC75, 1 *μ*l 150 *μ*M dC15, or 1 *μ*l 750 *μ*M dC5) and incubated 1 min at 22°C, and were supplemented with 1 *μ*l 0 or 30 µM Rho and incubated 5 min at 22°C. The resulting complexes were bead-immobilized by adding 1 *μ*l on streptavidin-coated agarose beads (Pierce Streptavidin Agarose; ThermoFisher; pre-washed with transcription buffer), incubating 10 min at 22°C, and removing unbound complex by washing with 3 × 100 *μ*l of transcription buffer. The resulting immobilized complexes were supplemented with 1 *μ*l 0 or 1 mM ATP (>99% purity; ThermoFisher) in transcription buffer and incubated 5 min at 37°C (to enable termination), were supplemented with 1 *μ*l 400 µM UTP, 400 µM CTP, and 1 mM 3’-d-GTP [>99% purity for UTP and CTP (ThermoFisher); ≥95% purity for 3’-dGTP (Jena Bioscience)] and incubated 2 min at 37°C (to “chase,” and thereby to stabilize, non-terminated complexes), and supernatants and beads were separated by centrifugation 1 min at 2,000xg at 22°C. Supernatant aliquots (10 µl) were removed and mixed with equal volumes of 2x stop buffer (90% formamide, 50 mM EDTA, 0.01% bromophenol blue, and 0.01% xylene cyanole) at 22°C, and pellets (beads) were washed with 60 µl 1x stop buffer (45% formamide, 25 mM EDTA, 0.005% bromophenol blue, and 0.005 xylene cyanole) at 22°C. Reaction products in supernatant and pellet factions were resolved by electrophoresis on 7 M urea, 20% polyacrylamide gels (19:1 acrylamide: bis-acrylamide; *7*), were detected using storage-phosphor imaging (Typhoon PhosphorImager, GE Healthcare), and were quantified using ImageQuant 5.2 (GE Healthcare).

### Scaffold assay for Rho-dependent transcription termination: RNA extension

Reaction mixtures) containing 0.5 µM RNAP core enzyme and 0.5 µM partial; nucleic-acid scaffold comprising template-strand oligodeoxyribonucleotide and 5’-^32^P-labelled RNA-product oligoribonucleotide (0.6 Bq/fmol) in 10 *μ*l transcription buffer were incubated 10 min at 22°C, were supplemented with 1 *μ*l 6 µM nontemplate-strand oligodeoxyribonucleotide and incubated 5 min at 22°C, were supplemented with 1 *μ*l 0 or 30 µM NusG derivative (NusG or NusG-N) and incubated 5 min at 22°C, were supplemented with PBS-ligand oligoribonucleotide (1 *μ*l 0 or 50 *μ*M λtR1 *rut*, 1 *μ*l 50 *μ*M dC75, 1 *μ*l 150 *μ*M dC15, or 1 *μ*l 750 *μ*M dC5) and incubated 1 min at 22°C, and were supplemented with 1 *μ*l 0 or 30 µM Rho and incubated 5 min at 22°C. The resulting reaction mixtures then were supplemented with 1 *μ*l 0 or 1 mM ATP (>99% purity; ThermoFisher) in transcription buffer and incubated 5 min at 37°C (to enable termination), were supplemented with 1 *μ*l mM 400 µM UTP, 400 µM CTP, and 400 µM 3’-dGTP [>99% purity for UTP and CTP (ThermoFisher); ≥95% purity for 3’-dGTP (Jena Bioscience)] and incubated 2 min at 37°C (to “chase” non-terminated complexes), and were mixed with 10 *μ*l 2x stop buffer at 22°C. Reaction products were resolved by electrophoresis on 7 M urea, 20% polyacrylamide gels (19:1 acrylamide: bis-acrylamide; *7*), were detected using storage-phosphor imaging (Typhoon PhosphorImager, GE Healthcare), and were quantified using ImageQuant 5.2 (GE Healthcare).

### Cryo-EM structure determination: sample preparation

Reaction mixtures containing 4 µM RNAP core enzyme and 5 *μ*M nucleic-acid scaffold comprising nontemplate-strand DNA, template-strand DNA and RNA product in 100 *μ*l transcription buffer were incubated 10 min at 22°C, were supplemented with 25 *μ*l 30 µM NusG derivative and incubated 5 min at 22°C, were supplemented with 3 *μ*l 0 or 1 mM PBS-ligand oligoribonucleotide (λtR1 *rut* or dC75) and incubated 1 min at 22°C, and were supplemented with 50 *μ*l 60 µM Rho and 2 µl 24 mM ADP-BeF_3_ and incubated 5 min at 22°C. The resulting complexes were concentrated to 35 µl (∼12 *μ*M final concentration) by centrifugation 12 min at 20,000xg at 4°C through pre-chilled Amicon Ultra-0.5 30K MWCO concentrators (EMD Millipore), supplemented with 4 µl of ice-cold 80 mM CHAPSO (Hampton Research), and kept on ice until applied to EM grids.

EM grids were prepared using a Vitrobot Mark IV autoplunger (FEI/ThermoFisher), with the environmental chamber set to 22°C and 100% relative humidity. Samples (3 μl) were applied to Quantifoil 1.2/1.3 Cu 300 holey-carbon grids (Quantifoil) glow-discharged 60 s using a PELCO glow-discharge system (Ted Pella); grids were blotted with #595 filter paper (Ted Pella) for 7-8 s at 22°C, flash-frozen by plunging in a liquid ethane cooled with liquid N_2_, and stored in liquid N_2_.

### Cryo-EM structure determination: data collection and data reduction: λtR1-NusG-Rho-TEC (n = 6)

Cryo-EM data for λtR1-NusG-Rho-TEC (n = 6) were collected at the National Center for CryoEM Access and Training (NCCAT), using a 300 kV Krios Titan (FEI/ThermoFisher) electron microscope equipped with a Gatan K2 Summit direct electron detector (Gatan), Leginon software (*8*), a nominal magnification of 81,000x, a calibrated pixel size of 1.069 Å/pixel, and a dose rate of 35 electrons/Å^2^/s. Movies were recorded at 30 ms/frame for 1.5 s (50 frames), resulting in a total radiation dose of 53 electrons/Å^2^. Defocus range was varied from -1.25 to -2.5. A total of 15,559 micrographs were recorded from one grid over three days. Micrographs were gain-normalized and defect-corrected.

Data were processed as summarized in Fig. S3. Data processing was performed using a Tensor TS4 Linux GPU workstation with four GTX 1080 Ti graphic cards (NVIDIA). Dose weighting and motion correction (3×3 tiles; b-factor = 150) were performed using Motioncor2 (9). CTF estimation was performed using CTFFIND-4.1 (*10*). Subsequent image processing was performed using Relion 3.0 (*11*). Automatic particle picking with Laplacian-of-Gaussian filtering yielded an initial set of 6,257,721 particles. Particles were extracted into 512×512 pixel boxes and subjected to rounds of reference-free 2D classification and removal of poorly populated classes, yielding a selected set of 69,316 particles. The selected set was 3D-classified with C1 symmetry, using a 3D template created *ab initio* using 3D_initial_model under Relion 3.0. Classes that exhibited strong, well-defined density for both Rho and the TEC were selected, 36,044 particles for the selected classes were combined, were 3D auto-refined using a mask with diameter of 200 Å, and were subjected to masked multi-body refinement using the Rho and TEC masks, yielding a reconstruction with a global resolution that reached 4.3 Å, as determined from gold-standard Fourier shell correlation (Fig S3D; Table S1). The reconstruction showed clear density for Rho, NusG, TEC, PBS-ligand RNA, SBS-ligand RNA, and spacer RNA (Fig S3E-I).

The initial atomic model for λtR1-NusG-Rho-TEC (n = 6) was built by docking of (i) the Rho hexamer, one copy of NusG-C, and Mg-ADP-BeF_3_ from a crystal structure of *E. coli* Rho hexamer interacting with six copies of *E. coli* NusG-C and Mg-ADP-BeF_3_ (PDB 6DUQ; *5*), (ii) SBS-ligand RNA from a crystal structure of a *E. coli* Rho hexamer interacting with SBS-ligand RNA and Mg-ADP-BeF_3_ (PDB 5JJI; *12*), and (iii) NusG-N, NusG linker, and the TEC from a cryo-EM structure of a complex of *E. coli* NusG bound to an *E. coli* TEC (PDB 6ZTJ; *13*), using Chimera (*14)*. PBS-ligand RNA, spacer RNA, and the double-stranded DNA upstream of the TEC were built manually using Chimera and Coot (*14-15*). For the Rho and C-termini (residues 418-420), the NusG N- and C-termini (residues 1-5 and 181-182), RNAP *β*’ N- and C-termini and trigger loop (residues 1 and 934-947, 1127-1135, and 1374-1408), RNAP *β* N- and C-termini (residues 1 and 1342-1343), RNAP *α*^I^ and *α*^II^ N-termini, activation-target loops, and C-terminal domains (residues 1-4, 159-164, and 234-329), and RNAP *ω* N- and C-termini (residues 1-2 and 75-92), density was absent, suggesting high segmental flexibility; these segments were not fitted.

Rigid-body refinement of TEC and Rho against the map was performed using real_space_refine under Phenix (*16*). PBS-ligand RNA, SBS-ligand RNA, spacer RNA, NusG-N (residues 45-62), NusG-C (residues 130-180), the NusG linker (residues 104-129), RNAP *β*’ ZBD (residues 62-95), RNAP *β* FTH (residues 890-912), and double-stranded DNA upstream of the TEC were subjected to iterative cycles of model building and refinement in Coot (*15*).

The final density maps for λtR1-NusG-Rho-TEC (n = 6) were deposited in the EMDB with accession codes EMD-27930 (map, processed by focused refinement, for TEC, NusG, and spacer RNA), EMD-27928 (map, processed by focused refinement, for Rho, PBS-ligand RNA, SBS-ligand RNA, and Mg-ADP-BeF_3_), and EMD-27929 (lower-resolution map of full complex, without focused refinement to improve resolution of components), and the final atomic coordinates were deposited in the PDB with accession codes PDB 8E6X (coordinates for TEC, NusG, SBS RNA, and spacer RNA) and PDB 8E6W (coordinates for Rho, PBS-ligand RNA, and Mg-ADP-BeF_3_) (Table S1).

### Cryo-EM structure determination: data collection and data reduction: dC75-NusG-Rho-TEC (n = 6)

Cryo-EM data for dC75-NusG-Rho-TEC (n = 6;) were collected at NCCAT, using a 300 kV Krios Titan (FEI/ThermoFisher) electron microscope equipped with a Gatan K2 Summit direct electron detector (Gatan) and Leginon software (*8*), using procedures as in the preceding section. A total of 16,394 micrographs were recorded from one grid over three days. Micrographs were gain-normalized and defect-corrected.

Data were processed as summarized in Fig. S4, using procedures as in the preceding section. Data processing yielded a reconstruction at 4.7 Å resolution (Fig. S4D; Table S1). Atomic models were built and refined as in the preceding section.

The final density maps of dC75-NusG-Rho-TEC (n = 6) were deposited in the EMDB with accession codes EMD-27931 (map, processed by focused refinement, for TEC, NusG, and spacer RNA), EMD-27932 (map, processed by focused refinement, for Rho, PBS-ligand RNA, SBS-ligand RNA, and Mg-ADP-BeF_3_), and EMD-27933 (lower-resolution map of full complex, without focused refinement to improve resolution of components), and the final atomic coordinates were deposited in the PDB with accession codes PDB 8E6Z (coordinates for TEC, NusG, SBS RNA, and spacer RNA) and PDB 8E70 (coordinates for Rho, PBS-ligand RNA, and Mg-ADP-BeF_3_) (Table S1).

### Cryo-EM structure determination: data collection and data reduction: NusG-Rho-TEC (n = 6. 7, and 8)

Cryo-EM data for NusG-Rho-TEC (n = 6) and NusG-Rho-TEC (n = 7) were collected at the Rutgers CryoEM and Nanoimaging Facility, using a 200 kV Talos Arctica (FEI/ThermoFisher) electron microscope equipped with a K2 Summit and Bioquantum energy filter (Gatan), SerialEM software (*17*), a nominal magnification of 130,000x, a calibrated pixel size of 1.038 Å/pixel, and a dose rate of 5 electrons/pixel/s, movies at 200 ms/frame for 6 s (30 frames; total radiation dose of 28 electrons/Å^2^), and a defocus range of -1.25 µm to -2.5 µm. A total of 2,784 micrographs were recorded for NusG-Rho-TEC (n = 6) from one grid over one day, and a total of 8,528 micrographs were recorded for NusG-Rho-TEC (n = 7) from one grid over two days. Micrographs were gain-normalized and defect-corrected.

Cryo-EM data for NusG-Rho-TEC (n = 8) were collected at the Stanford-SLAC Cryo-EM Center (SLAC), using a 300 kV Krios Titan (FEI/ThermoFisher) electron microscope equipped with a Gatan K2 Summit direct electron detector (Gatan), SerialEM software, a nominal magnification of 81,000x, a calibrated pixel size of 1.0 Å/pixel, a dose rate of 55 electrons/Å^2^/s, movies at 35 ms/frame for 1.1 s (30 frames;, total radiation dose of 50 electrons/Å2), and a defocus range of -1.25 μm to -2.5 μm. A total of 15,340 microphotographs were recorded from one grid over three days. Micrographs were gain-normalized and defect-corrected.

Data were processed as summarized in Fig. S5, using procedures as in the preceding two sections. Data processing yielded reconstructions at 6.5, 4.4, and 4.2 Å resolution for NusG-Rho-TEC (n = 6), NusG-Rho-TEC (n = 7), and NusG-Rho-TEC (n = 8), respectively (Fig. S5D; Table S1). Atomic models were built and refined as in the preceding two sections.

The final density maps of NusG-Rho-TEC (n = 6) were deposited in the EMDB with accession codes EMD-27864 (map, processed by focused refinement, for TEC, NusG, and spacer RNA), EMD-27865 (map, processed by focused refinement, for Rho, PBS-ligand RNA, SBS-ligand RNA, and Mg-ADP-BeF_3_), and EMD-27897 (lower-resolution map of full complex, without focused refinement to improve resolution of components), and the final atomic coordinates were deposited in the PDB with accession codes PDB 8E3F (coordinates for TEC, NusG, SBS RNA, and spacer RNA) and PDB 8E3H (coordinates for Rho, PBS-ligand RNA, and Mg-ADP-BeF_3_) (Table S1).

The final density maps of NusG-Rho-TEC (n = 7) were deposited in the EMDB with accession codes EMD-27913 (map, processed by focused refinement, for TEC, NusG, and spacer RNA), EMD-27914 (map, processed by focused refinement, for Rho, PBS-ligand RNA, SBS-ligand RNA, and Mg-ADP-BeF_3_), and EMD-27915 (lower-resolution map of full complex, without focused refinement to improve resolution of components), and the final atomic coordinates were deposited in the PDB with accession codes PDB 8E5K (coordinates for TEC, NusG, SBS RNA, and spacer RNA) and PDB 8E5L (coordinates for Rho, PBS-ligand RNA, and Mg-ADP-BeF_3_) (Table S1).

The final density maps of NusG-Rho-TEC (n = 8) were deposited in the EMDB with accession codes EMD-27916 (map, processed by focused refinement, for TEC, NusG, and spacer RNA), EMD-27917 (map, processed by focused refinement, for Rho, PBS-ligand RNA, SBS-ligand RNA, and Mg-ADP-BeF_3_), and EMD-27918 (lower-resolution map of full complex, without focused refinement to improve resolution of components), and the final atomic coordinates were deposited in the PDB with accession codes PDB 8E5O (coordinates for TEC, NusG, SBS RNA, and spacer RNA) and PDB 8E5P (coordinates for Rho, PBS-ligand RNA, and Mg-ADP-BeF_3_) (Table S1).

## Supplementary Figure Legends

**Fig. S1.**
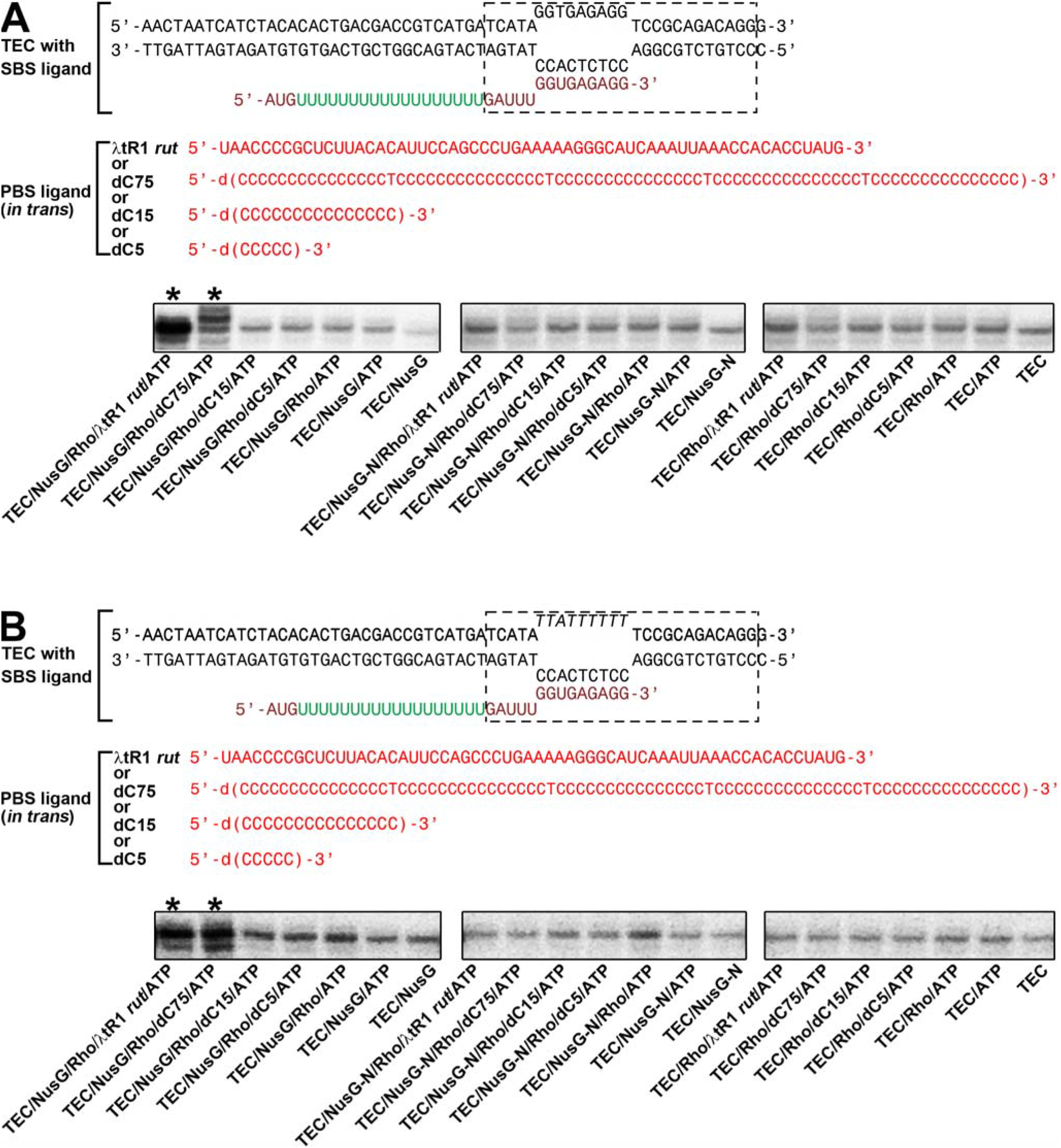
Scaffold assay for Rho-dependent termination: additional data. **(A)** Top, **s**ynthetic nucleic-acid scaffolds containing complementary nontemplate and template DNA strands (colors as in Fig. 1B). Bottom, RNA-release data assessing Rho-dependent termination in complexes containing NusG (left), NusG-N (center), or no NusG (right). Asterisks, released RNA products that exhibit increased levels relative to control reactions lacking Rho (lanes 6 and 7 of each subpanel). Termination is detected by an increase in released RNA products relative to control reactions lacking Rho. **(B)** As in (A), except that synthetic nucleic-acid scaffolds contain noncomplementary nontemplate and template DNA strands.

**Fig. S2.**
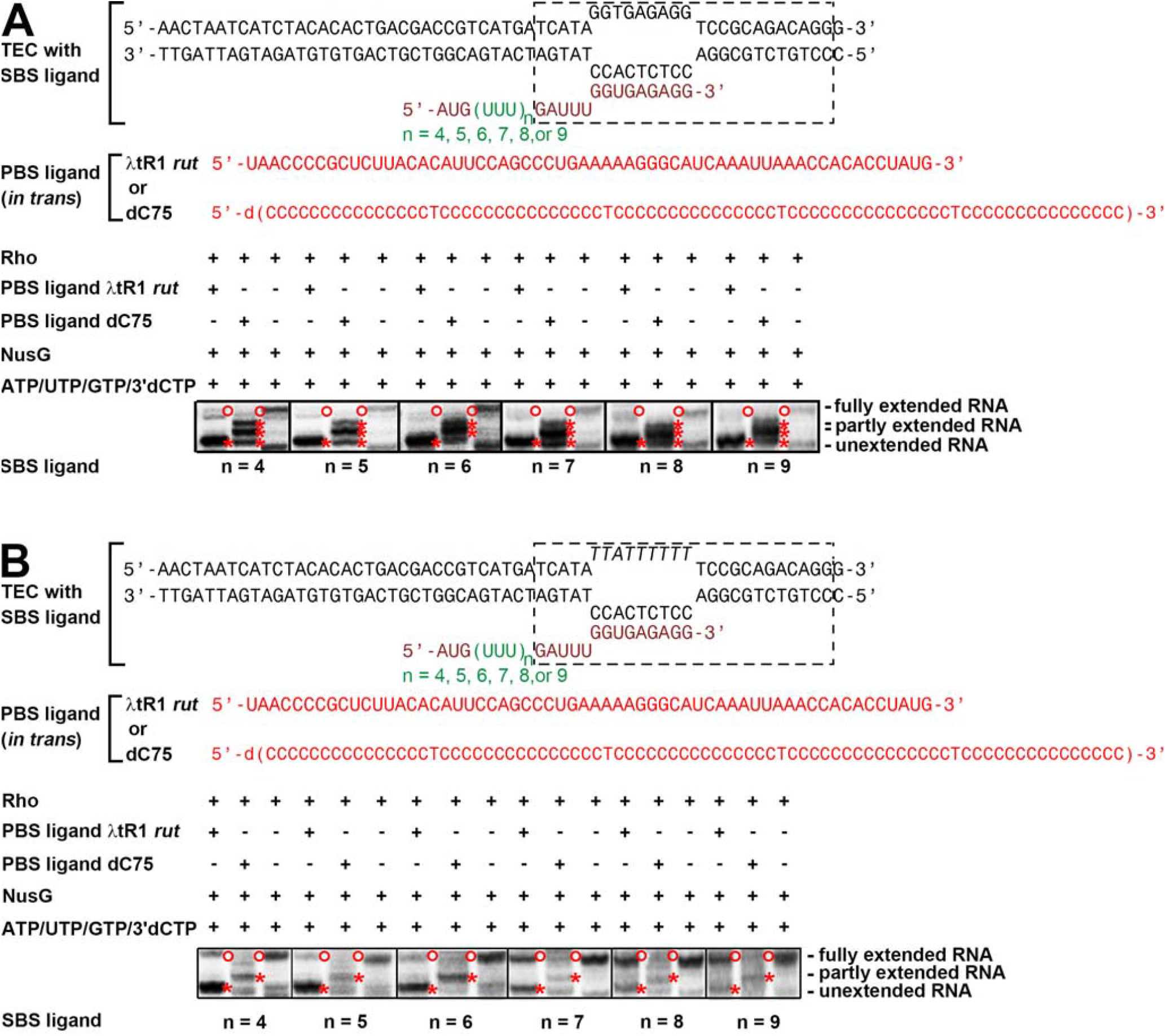
Scaffold assay for Rho-dependent termination: additional data. **(A)** Top, **s**ynthetic nucleic-acid scaffolds containing complementary nontemplate and template DNA strands (colors as in Figs. 1B, S1). Bottom, RNA-extension data assessing Rho-dependent termination in complexes having λtR1 *rut* as PBS ligand (odd-numbered lanes) or dC75 as PBS ligand (even-numbered lanes) on scaffolds where spacer length, n, is 4, 5, 6, 7, 8 or 9 codons (first, second, third, fourth, fifth, and sixth subpanels). Red asterisks, RNA products that exhibit increased levels compared to control reactions without PBS ligand; open circles, RNA products that exhibit decreased levels relative to control reactions without PBS ligand. Termination is detected by an increase in unextended and partly extended RNA products (lower and central bands), and a decrease in fully extended RNA products (top bands). **(B)** As in (A), except that **s**ynthetic nucleic-acid scaffolds contain noncomplementary nontemplate and template DNA strands.

**Fig. S3.**
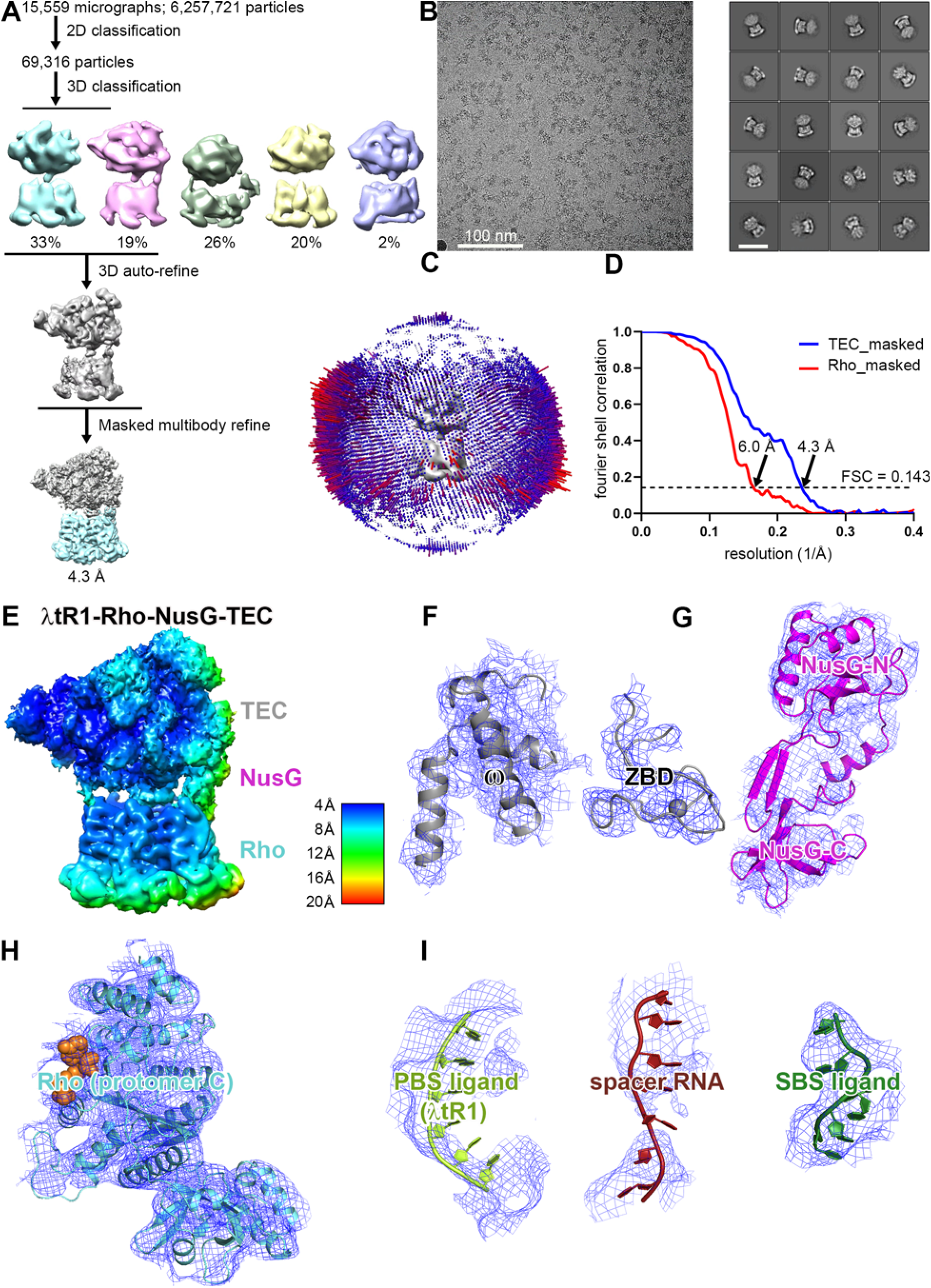
Structure determination: λtR1-NusG-Rho-TEC (n = 6) **(A)** Data processing scheme (Table S1). **(B)** Representative electron microphotograph and 2D class averages (50 nm scale bar in right subpanel). **(C)** Orientation distribution. **(D)** Fourier-shell correlation (FSC) plot. **(E)** EM density map colored by local resolution (view orientation as in Fig. 2A, left). **(F-I)** Representative EM density (blue mesh) and fits (ribbons) for RNAP regions that interact with Rho, NusG, Rho protomer C, PBS ligand interacting with Rho protomer C, spacer RNA, and SBS ligand.

**Fig. S4.**
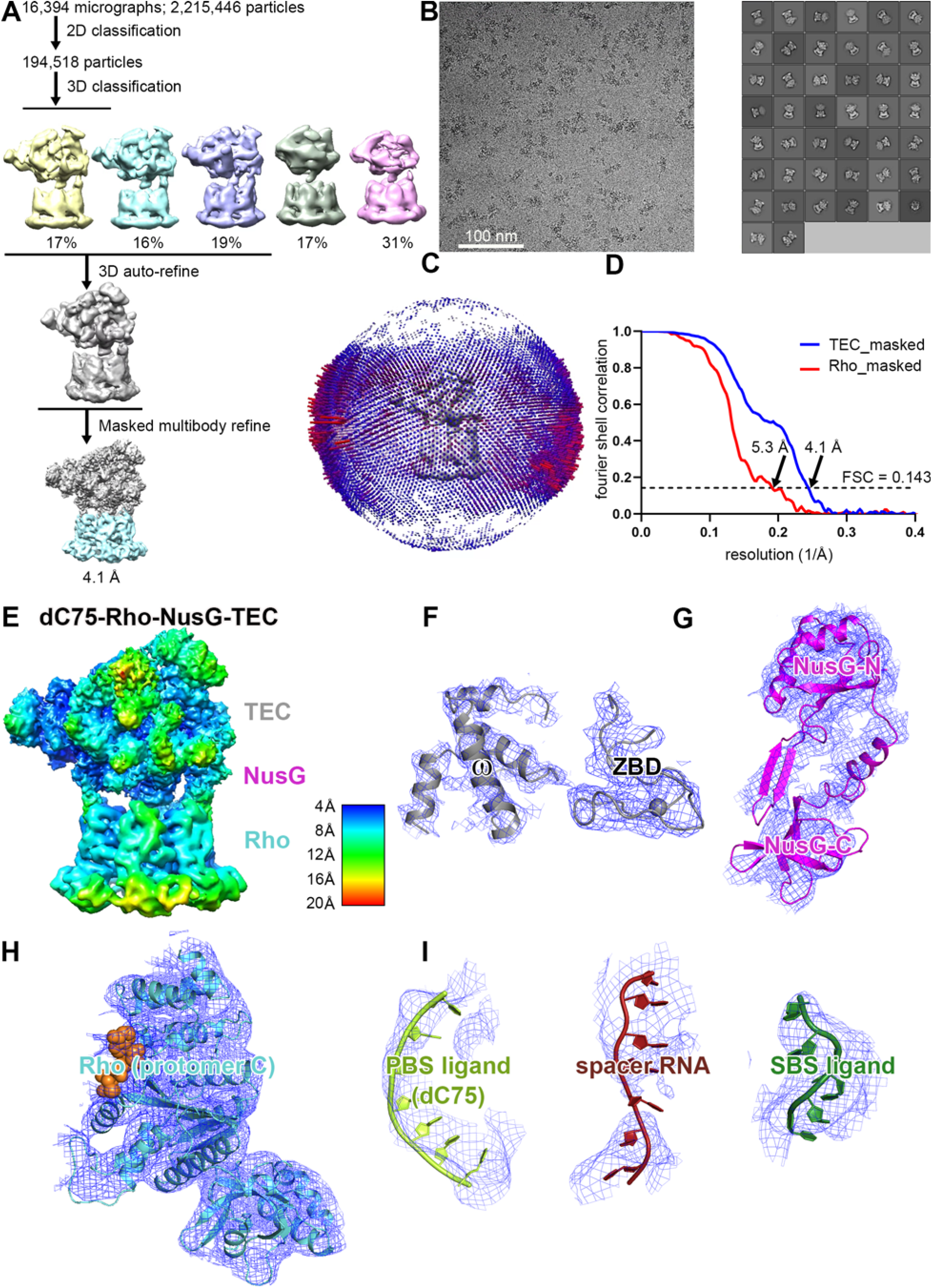
Structure determination: dC75-NusG-Rho-TEC (n = 6) **(A)** Data processing scheme (Table S1). **(B)** Representative electron microphotograph and 2D class averages (50 nm scale bar in right subpanel). **(C)** Orientation distribution. **(D)** Fourier-shell correlation (FSC) plot. **(E)** EM density map colored by local resolution (view orientation as in Fig. 2A, left). **(F-I)** Representative EM density (blue mesh) and fits (ribbons) for RNAP regions that interact with Rho, NusG, Rho protomer C, PBS ligand interacting with Rho protomer C, spacer RNA, and SBS ligand.

**Fig. S5.**
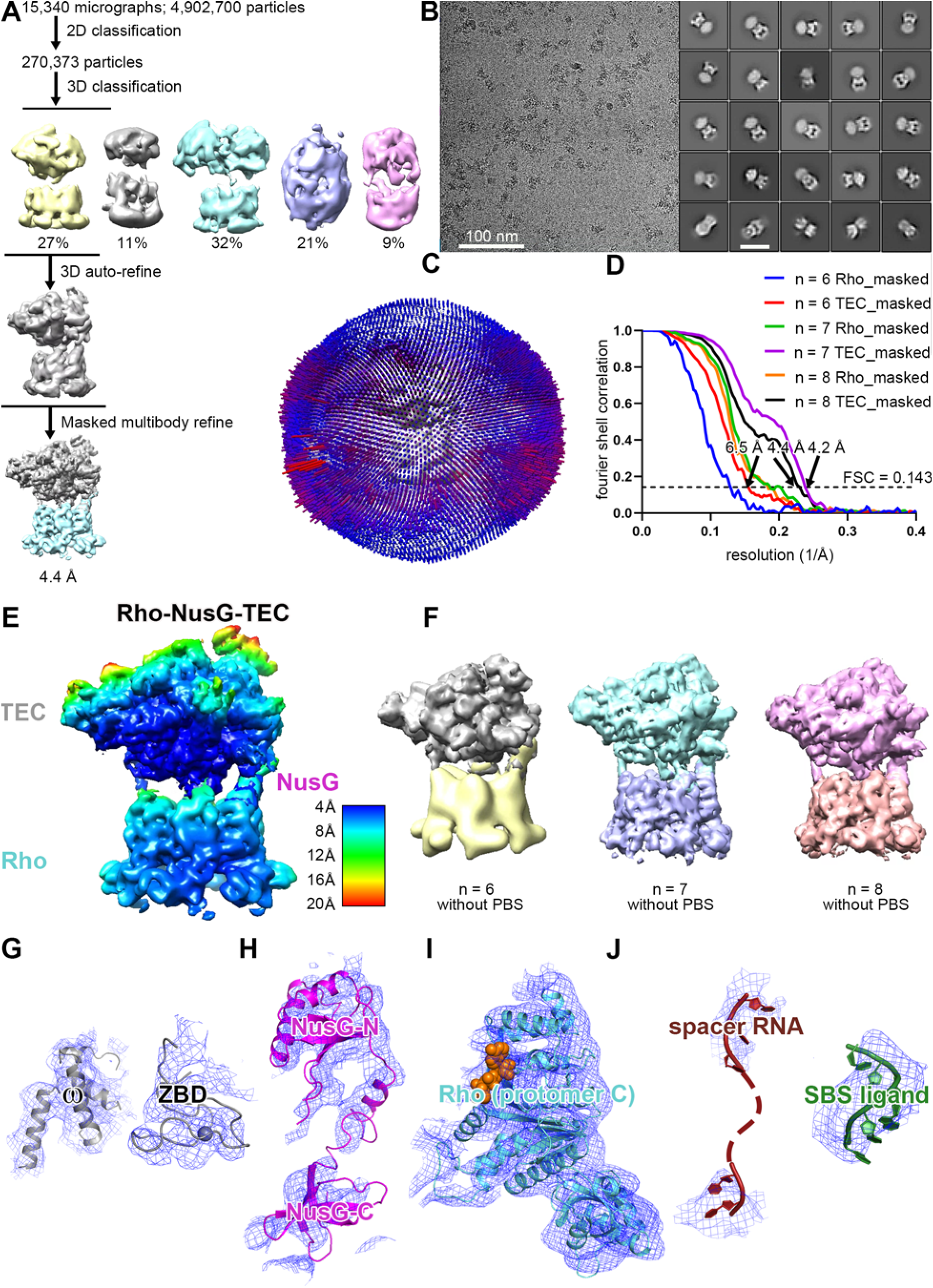
Structure determination: NusG-Rho-TEC (n = 6, 7, and 8) **(A)** Data processing scheme (n = 8). **(B)** Representative electron microphotograph and 2D class averages (n = 8; 50 nm scale bar in right subpanel). **(C)** Orientation distribution (n = 8). **(D)** Fourier-shell correlation (FSC) plot. **(E)** EM density map colored by local resolution (n = 8) (view orientation as in Fig. 2A, left). **(F)** EM density maps for NusG-Rho-TEC obtained using nucleic acid scaffolds with n = 6, 7, and 8 (view orientation as in Fig. 2A, left). **(G-J)** Representative EM density (blue mesh) and fits (ribbons) for RNAP regions that interact with Rho, NusG, Rho protomer C, spacer RNA, and SBS ligand.

**Fig. S6.**
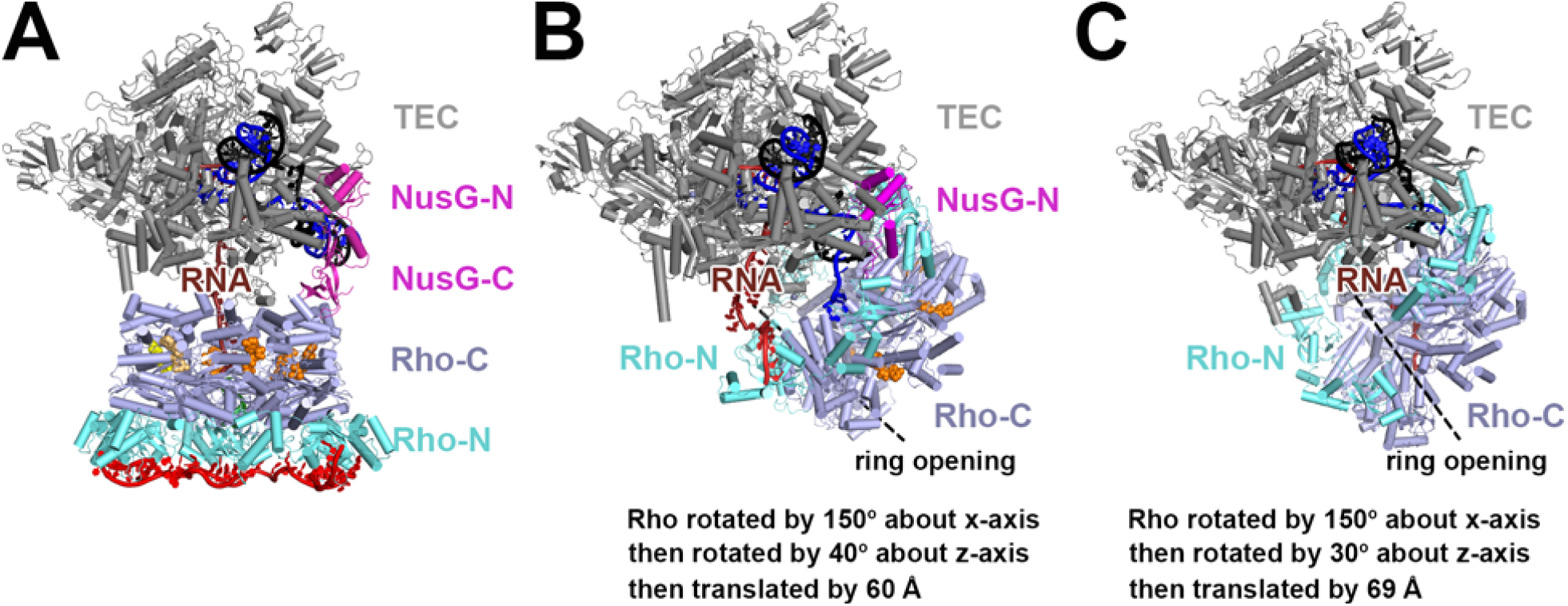
Comparison of structure of λtR1-NusG-Rho-TEC to structures of Rho-TEC complexes of *18* and *19*. **(A)** Structure of λtR1-NusG-Rho-TEC. Rho N-terminal domain (Rho-N); cyan, and Rho C-terminal domain (Rho-C), slate blue, are shown in different colors to highlight orientation of Rho domains relative to the TEC. View orientation and other colors as in Fig. 2A, left. **(B)-(C)** Structures of Rho-TEC complexes of *18* (B; PDB 6Z9R) and *19* (C; PDB 6XAS; NusA omitted for clarity). View orientation that superimposes TEC atoms in (B) and (C) on TEC atoms in (A). Colors as in (A). The ring opening of the open-ring Rho hexamer in structures of *18* and *19* is indicated by a dashed line, and rotations and translation that relate the orientation of Rho relative to the TEC in structures of *18* and *19* to those in (A) are summarized below.

**Fig. S7.**
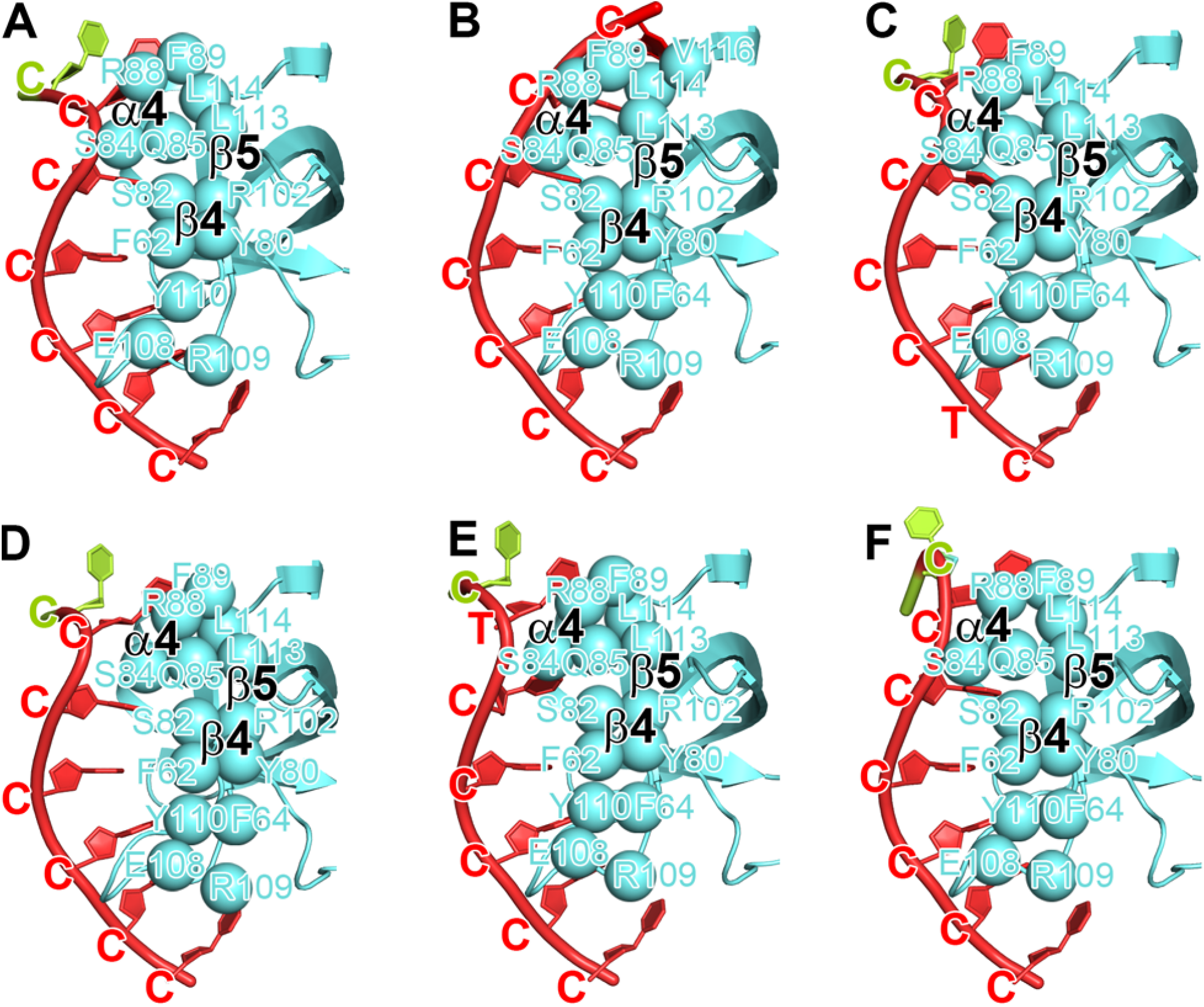
Protein-RNA interactions between PBS ligand and Rho PBS: dC75-NusG-Rho-TEC. Rho-(PBS ligand) interactions in dC75-NusG-Rho-TEC. View orientations and colors as in Fig. 3A, top right.

**Fig. S8.**
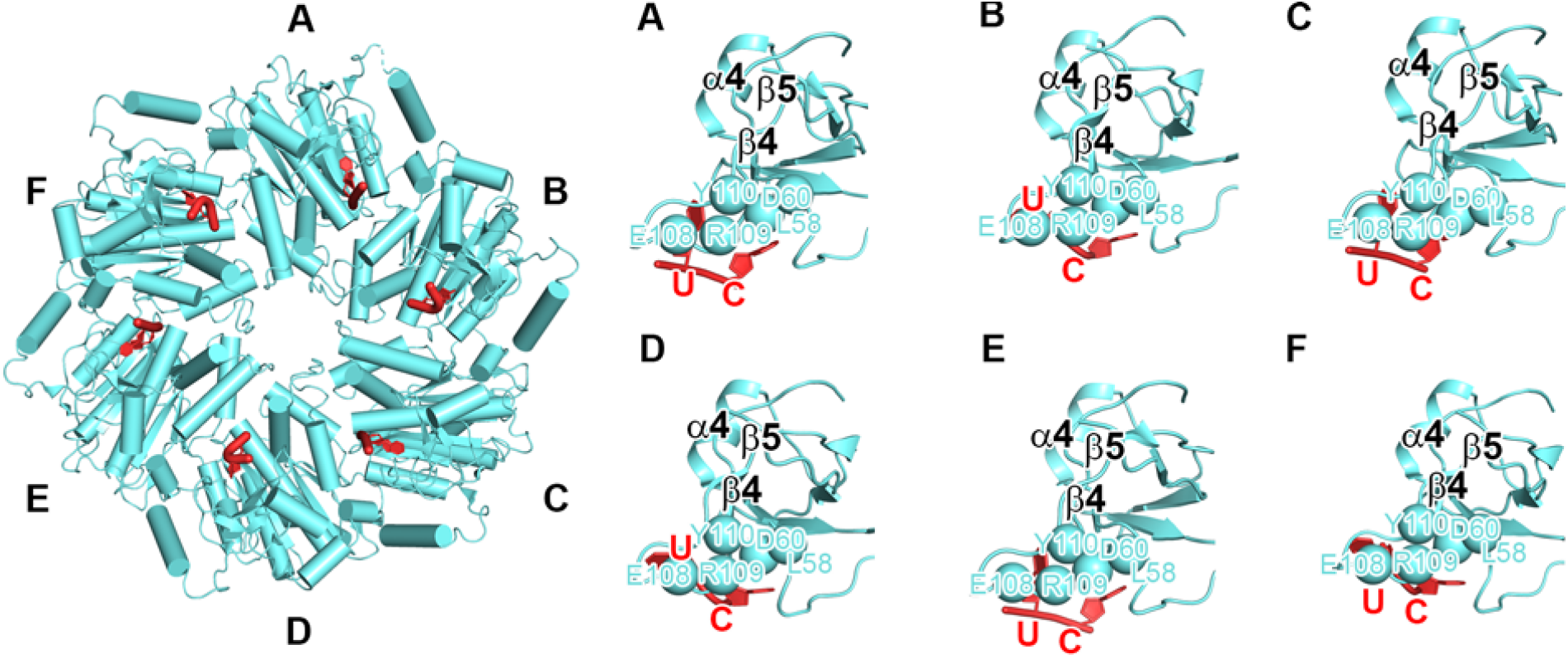
Protein-RNA interactions between PBS ligand and Rho PBS: crystal structure of Rho hexamer interacting with six copies of oligoribonucleotide in absence of NusG and TEC. Rho-(PBS ligand) interactions in crystal structure of Rho hexamer interacting with six copies of oligoribonucleotide in absence of NusG and TEC (*20*; PDB 2HT1). View orientations and colors as in Fig. 3A, top. Oligoribonucleotides non-specifically interacting with Rho-C omitted for clarity.

**Fig. S9.**
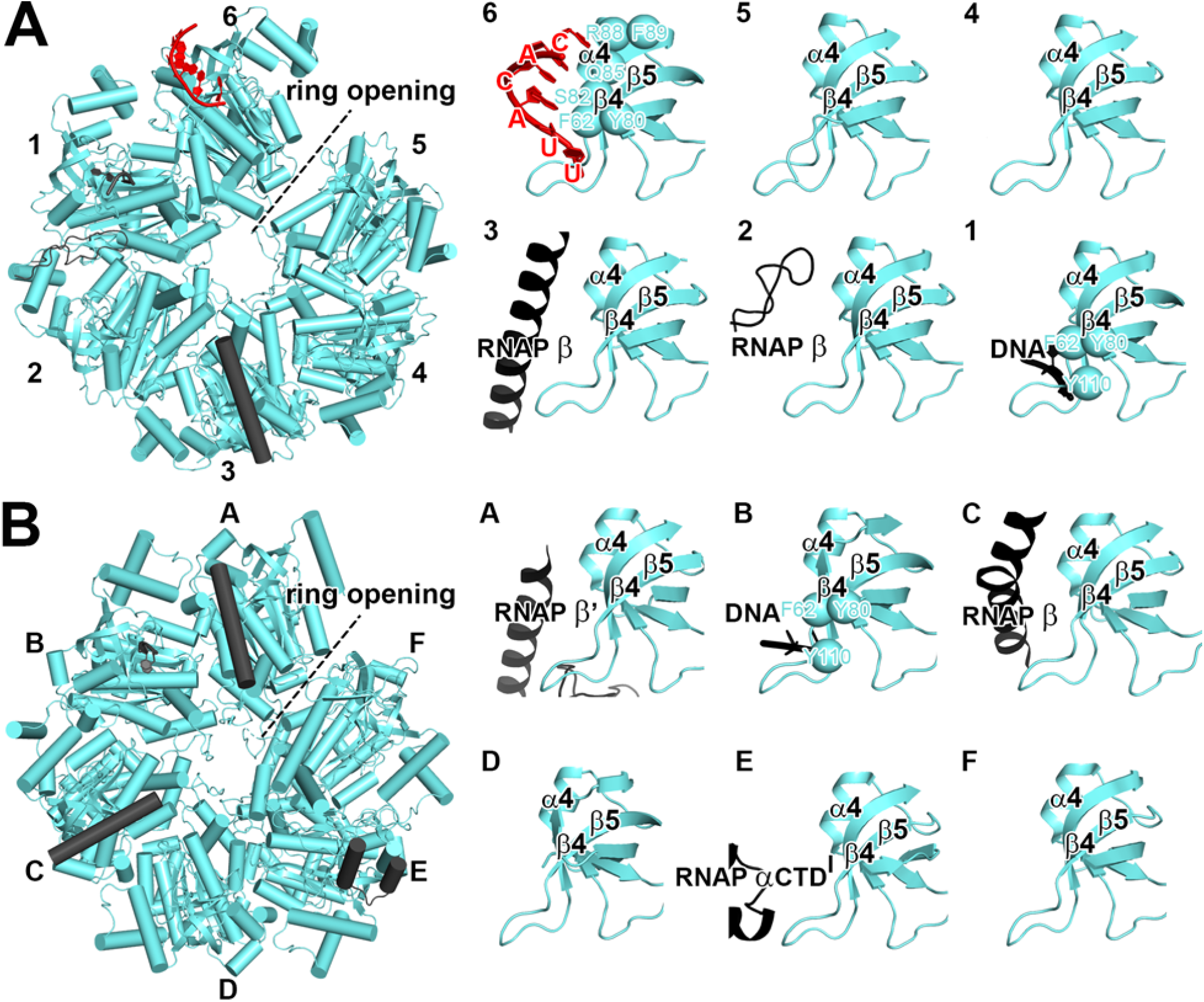
Protein-RNA interactions between PBS ligand and Rho PBS: structures of *18* and *19*. Rho-(PBS ligand) interactions in **s**tructures of *18* (A; PDB 6Z9R) and *19* (B; PDB 6XAS). Dashed black lines, ring openings in open-ring Rho hexamers in structures of *18* and *19*; black cylinders and black ribbons, RNAP or DNA structural elements of that occlude PBS-binding sites, preventing interaction with PBS ligand in structures of *18* and *19*. View orientations and other colors as in Fig. 3A, top.

**Fig. S10.**
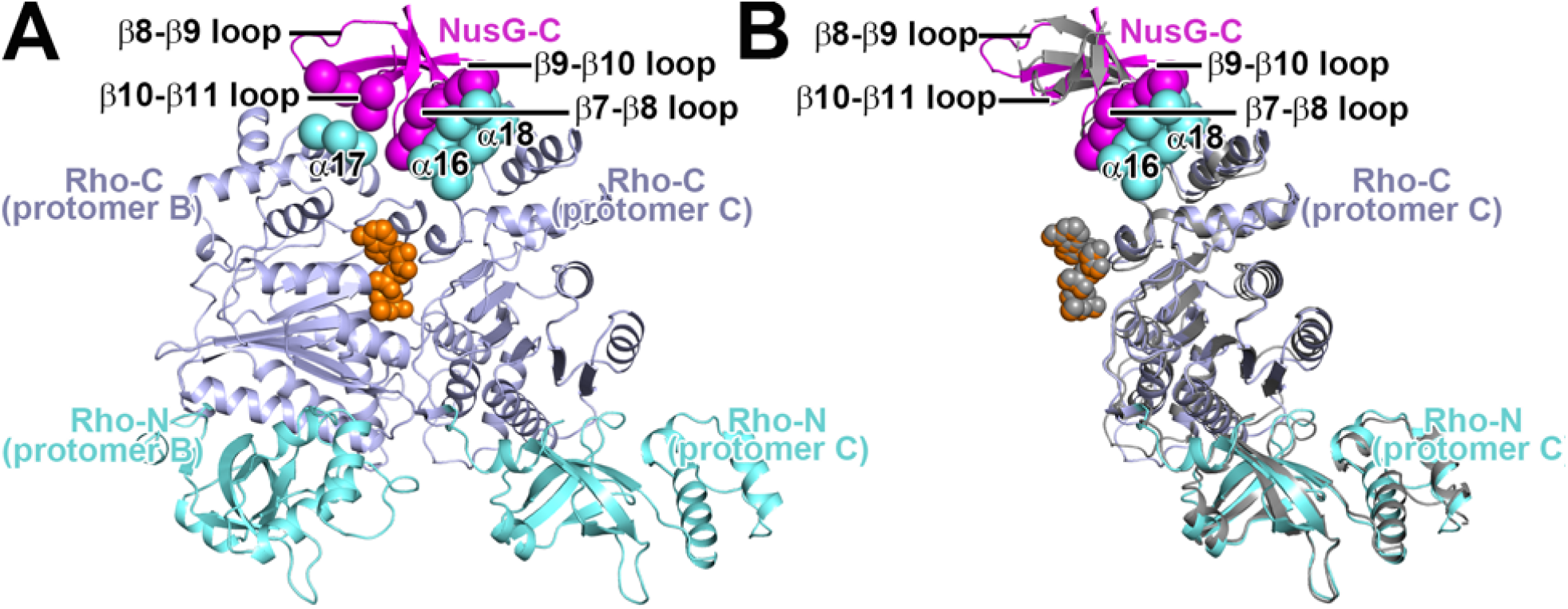
Protein-protein interactions between NusG-C and Rho. **(A)** Rho-(NusG-C) interactions in *λ*tR1-NusG-Rho-TEC. Colors as in Fig. S6. **(B)** Superimposition of NusG-C and Rho protomer C of λtR1-NusG-Rho-TEC (colored as in (A) on structure of one NusG-C and one Rho protomer in crystal structure of Rho hexamer interacting with six copies of NusG-C in absence of TEC (*21*; PDB 6DUQ; gray). Dashed grey ribbon, disordered segment of NusG-C *β*8-*β*9 loop in crystal structure.

**Fig. S11.**
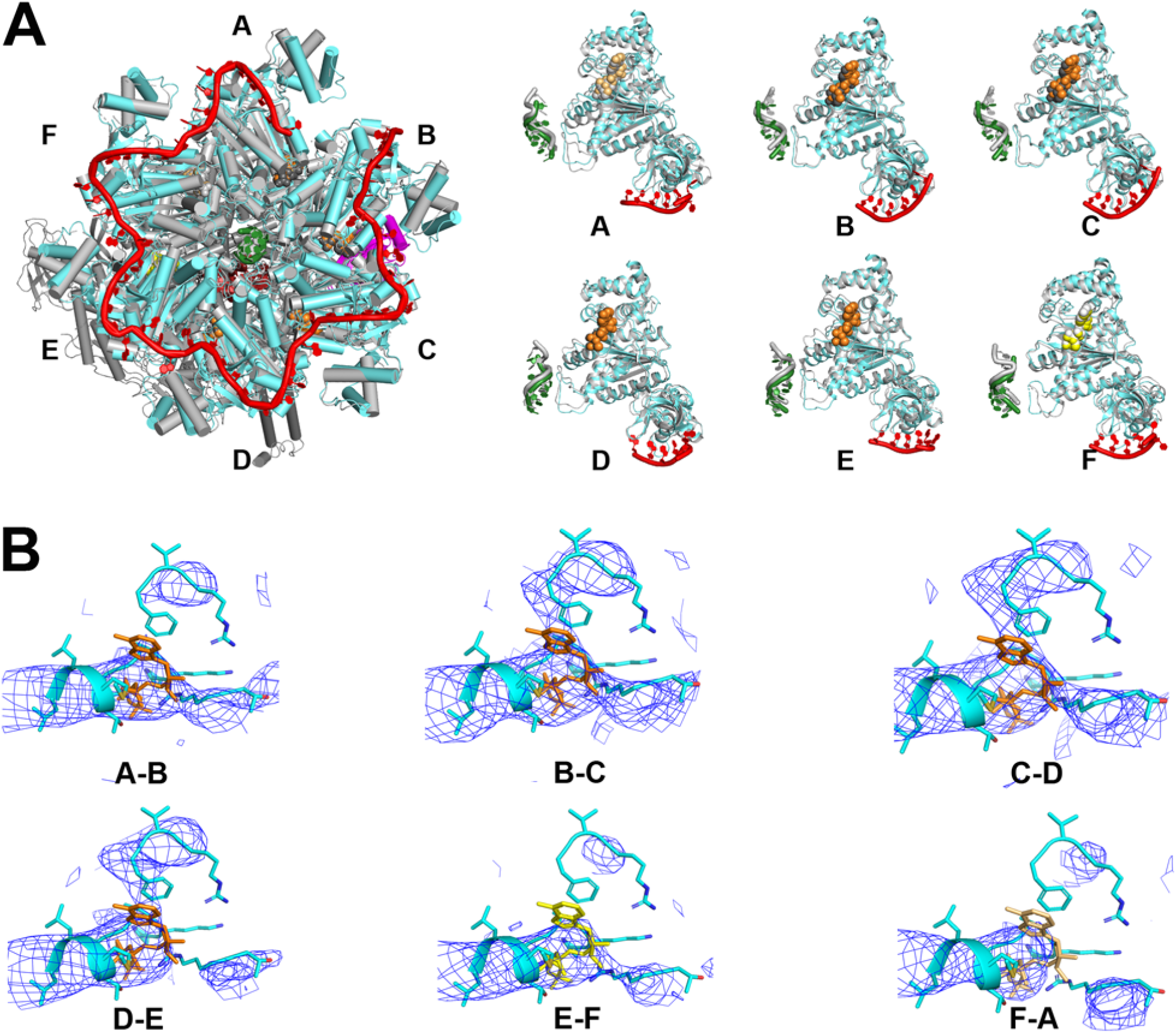
Motor states of Rho hexamer. **(A)** Left, superimposition of structure of λtR1-NusG-Rho-TEC (colored as in Fig. 2A, left) on crystal structure of Rho hexamer interacting with SBS ligand and Mg-ADP-BeF_3_ in absence of NusG and TEC (*22*; PDB 5JJI; colored gray). View orientation as in Fig. 2A, left. TEC omitted for clarity. Right, **s**uperimposition of Rho protomers A-F interacting with PBS ligand, SBS ligand, and Mg-ADP-BeF_3_ in structure of λtR1-NusG-Rho-TEC (colored as in A) on crystal structure of Rho protomers A-F interacting with SBS ligand and Mg-ADP-BeF_3_ (*22*; PDB 5JJI; gray). **(B)** Occupancy and order of ATP-binding sites of λtR1-NusG-Rho-TEC. Figure presents EM density (blue mesh) and fit (cyan for Rho; orange, light orange, and yellow for Mg-ADP-BeF_3_ at high, low, and very low occupancies, respectively) for ATP binding sites between Rho protomers A and B, B and C, C and D, D and E, E and F, and F and A.

**Fig. S12.**
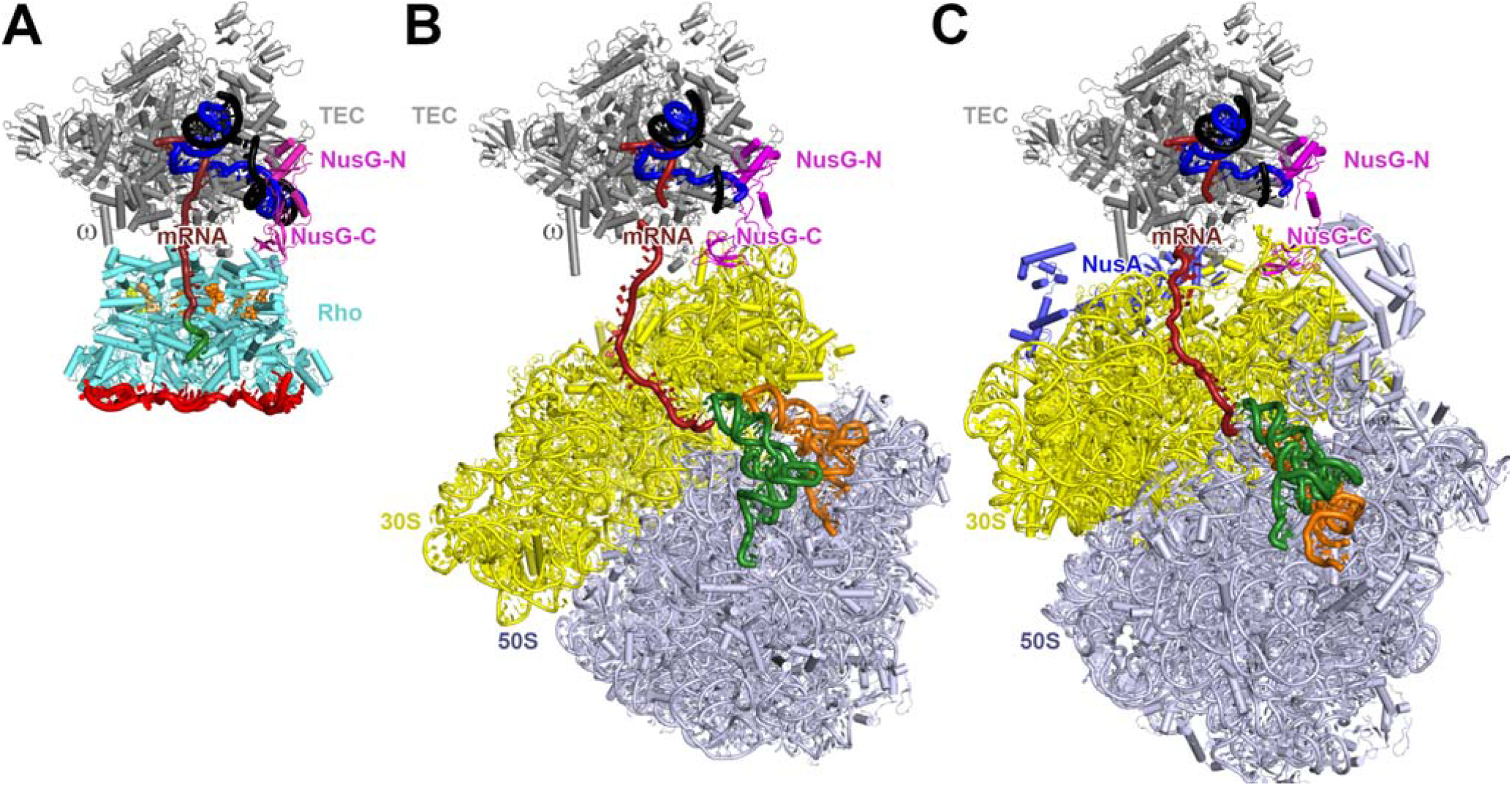
Comparison of structure of λtR1-NusG-Rho-TEC to structures of functional transcription-translation complexes NusG-TTC-B and NusA-NusG-TTC-B. **(A)** Structure of λtR1-NusG-Rho-TEC. View orientation and colors as in Fig. 2A, left. **(B)-(C)** Structures of NusG-TTC-B (B; *13, 23*; PDB 6XII) and NusA-NusG-TTC-B (C; *23*; PDB 6X7F). View orientation that superimposes TEC atoms in (B) and (C) on TEC atoms in (A). Ribosome 30S subunit, yellow; ribosome 50S subunit, gray; P- and E-site tRNAs bound to ribosome, green and orange; NusA, light blue. Other colors as in A.

**Table S1.** Cryo-EM structures: *λ*tR1-Rho-NusG-TEC (n = 6), dC75-Rho-NusG-TEC (n = 6), and Rho-NusG-TEC (n = 6, 7, and 8)

| complex | n (codons) | facility | particles | resolution | EMDB code<sup>a</sup> | PDB code<sup>b</sup> |
| --- | --- | --- | --- | --- | --- | --- |
| λtR1-Rho-NusG-TEC | 6 | NCCAT | 69,316 | 4.3 Å | EMD-27930<br>EMD-27928<br>EMD-27929 | 8E6X<br>8E6W |
| dC75-Rho-NusG-TEC | 6 | NCCAT | 194,518 | 4.1 Å | EMD-27931<br>EMD-27932<br>EMD-27933 | 8E6Z<br>8E70 |
| Rho-NusG-TEC | 6 | Rutgers | 15,936 | 6.5 Å | EMD-27864<br>EMD-27865<br>EMD-27897 | 8E3F<br>8E3H |
| Rho-NusG-TEC | 7 | Rutgers | 31,098 | 4.2 Å | EMD-27913<br>EMD-27914<br>EMD-27915 | 8E5K<br>8E5L |
| Rho-NusG-TEC | 8 | SLAC | 270,373 | 4.4 Å | EMD-27916<br>EMD-27917<br>EMD-27918 | 8E5O<br>8E5P |

<sup>a</sup>The first, second, and third EMDB codes for each complex are for (i) density map, processed by focused refinement, for TEC, NusG, and spacer RNA, (ii) density map, processed by focused refinement, for Rho, PBS-ligand RNA (where present), SBS-ligand RNA, and Mg-ADP-BeF<sub>3</sub>, and (iii) lower-resolution density map of full complex, without focused refinement to improve resolution of components.
<sup>b</sup>The first and second PDB codes for each complex are for (i) atomic coordinates for TEC, NusG, SBS-ligand RNA, and spacer RNA and (ii) atomic coordinates for Rho, PBS-ligand RNA (where present), and Mg-ADP-BeF<sub>3</sub>.

